# The Time Complexity of Self-Assembly

**DOI:** 10.1101/2021.04.01.437956

**Authors:** Florian M. Gartner, Isabella R. Graf, Erwin Frey

**Affiliations:** Arnold Sommerfeld Center for Theoretical Physics (ASC) and Center for NanoScience (CeNS), Department of Physics, Ludwig-Maximilians-Universität München, Theresienstraße 37, 80333 München, Germany

**Author notes:** Present address: Department of Physics, Yale University, New Haven, CT 06520, USA.

## Abstract

Time efficiency of self-assembly is crucial for many biological processes. Moreover, with the advances of nanotechnology, time efficiency in artificial self-assembly becomes ever more important. While structural determinants and the final assembly yield are increasingly well understood, kinetic aspects concerning the time efficiency, however, remain much more elusive. In computer science, the concept of *time complexity* is used to characterize the efficiency of an algorithm and describes how the algorithm’s runtime depends on the size of the input data. Here we characterize the time complexity of non-equilibrium self-assembly processes by exploring how the time required to realize a certain, substantial yield of a given target structure scales with its size. We identify distinct classes of assembly scenarios, i.e. ‘algorithms’ to accomplish this task, and show that they exhibit drastically different degrees of complexity. Our analysis enables us to identify optimal control strategies for non-equilibrium self-assembly processes. Furthermore, we suggest an efficient irreversible scheme for the artificial self-assembly of nanostructures, which complements the state-of-the-art approach using reversible binding reactions and requires no fine-tuning of binding energies.

## 1 Introduction

Time efficiency of self-assembly plays an important role in biology. For example, virus assembly must be fast in order to produce many virus particles before the infected cell is eliminated by the host’s immune system [14, 22, 37]. Moreover, as larger and ever more complex nanostructures are to be realized for technological or medical applications, time efficiency in artificial self-assembly becomes vital [7, 30]. Designing self-assembly schemes that are fast and resource efficient is, however, challenging. The task amounts to finding strategies that avoid the formation of large numbers of incompatible and incomplete fragments of the desired target structure. Such kinetic traps [8, 9, 15, 18, 32] arise even when all building blocks have a high binding specificity and erroneous binding is negligible, and they become more prominent with increasing structure size. Consequently, assembly time increases with structure size. But how exactly does the assembly time scale with the size of the target structure, and how does this scaling depend on the self-assembly scheme? What kinds of schemes optimize the assembly time? Answers to these questions will enable assembly strategies to be identified that are optimally suited for the production of large, functionally complex macromolecular structures via artificial self-assembly, a major goal in nanotechnology [5, 7, 24, 27, 30]. Here, we address these questions by studying the time complexity (as opposed to structural complexity [1, 6, 10, 29]) of four prototypical self-assembly scenarios, using scaling arguments and in-silico modelling of the stochastic dynamics (see Methods). Three of these scenarios have well-established realizations in biological and artificial self-assembly processes. The fourth strategy is a distinct idea conceptualized to achieve efficient self-assembly in a technological context by effectively regulating the supply of building blocks.

## 2 General model and self-assembly scenarios

To explore these questions in their simplest form, we consider an assembly process involving *N* identical copies of *S* different species of building blocks (monomers) and assume chemical reaction kinetics in a well-mixed fluid environment. As we expect the time efficiency of the assembly process to depend on the dimensionality of the structure, we investigate the assembly of linear polymers, two-dimensional sheets, and three-dimensional cubes of edge length *L* (size *S*) (Fig. 1). We assume the following reaction kinetics: Any two compatible monomers can bind at rate *μ*, forming a dimer that serves as a nucleus for further growth by sequential addition of monomers at rate *ν* per binding site. Following the assumptions of classical aggregation theory, we neglect interactions among polymers [3, 6]. Analyses of more complex reaction schemes including heterogeneous binding rates are discussed in the Supplementary Information (SI) and show that our conclusions are robust against model modifications. We will mainly consider irreversible processes, in which structures can only grow. To assess the relevance of reversible binding, we also discuss a scenario in which individual monomers may detach from the edges of incomplete structures at a finite detachment rate *δ_n_* that decreases exponentially with the number *n* of bonds that need to be broken: 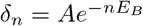 (Arrhenius’ law). Here *E_B_* is the binding energy per contact (bond) in units of *k_B_T* and the constant *A* can typically be assumed to be large relative to the rate of reactions [4, 26]. Once a structure contains all *S* species it is considered complete, and no further attachment or detachment processes occur (absorbing state). The yield of the assembly process is defined as the number of complete structures relative to their maximum possible number *N*.

**Figure 1.**
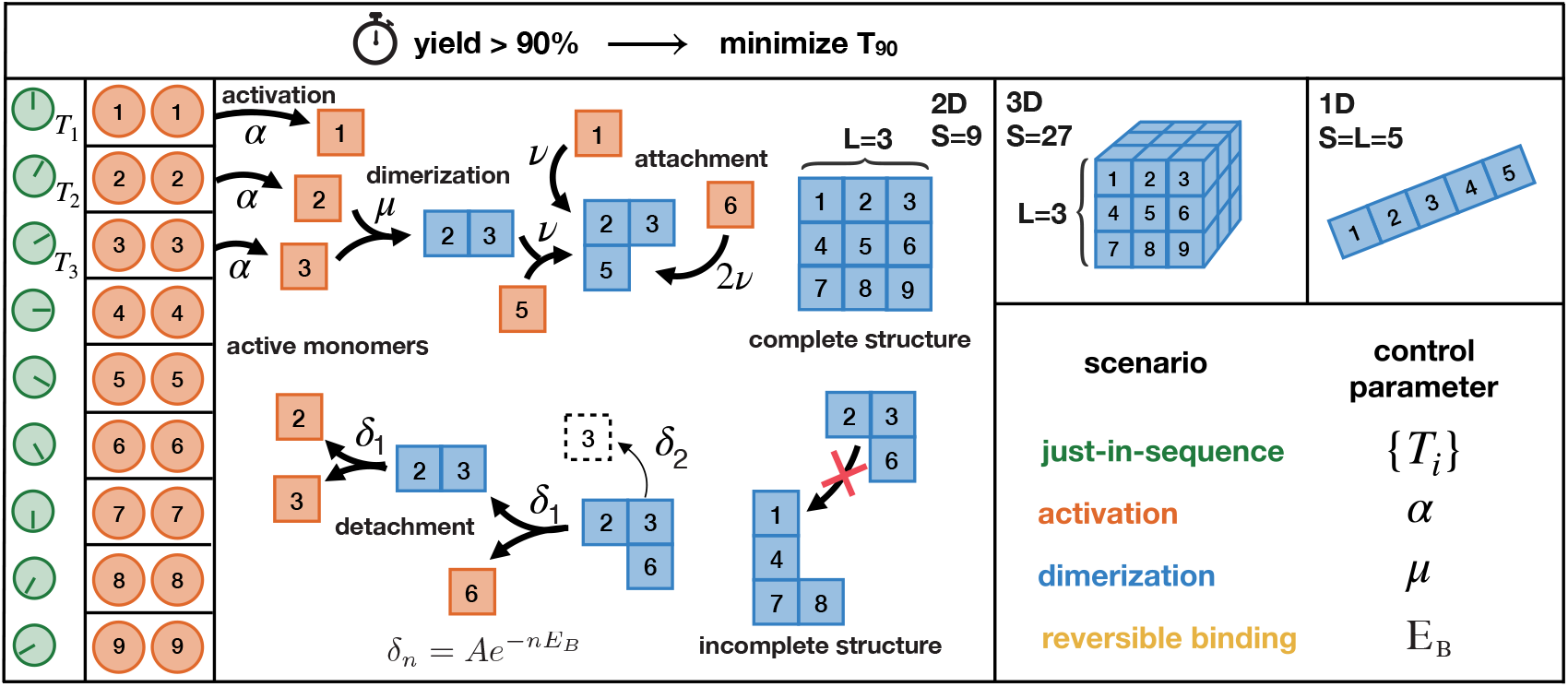
Schematic description of the model. *N* identical copies of *S* different species of monomers assemble into one- (1D), two- (2D) or three-dimensional (3D) heterogeneous structures of edge length *L* (only the 2D case is illustrated explicitly). A constant influx of monomers of species *i* takes place during the time interval 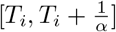 with net influx rate *Nα*. Once added to the system (activated), monomers start to self-assemble. A monomer of a bulk species has 2 (1D), 4 (2D) or 6 (3D) possible binding partners as illustrated in the figure. Any two fitting monomers can dimerize with rate *μ*. Subsequent to dimerization, structures grow by attachment of single monomers with rate *ν* per binding site. Furthermore, monomers can detach from a cluster with rate 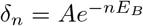, where *n* is the number of bonds that need to be broken and *E_B_* the binding energy per bond. We set *A* = 10^18^ *Cν*, with *C* = *N/V* denoting the concentration of monomers per species. Our aim is to minimize the assembly time *T*_90_ when 90% of all resources are assembled into complete structures. To this end, we control particular elements of the assembly process (control parameters) and distinguish four scenarios which are defined through the respective control parameter(s). The other parameters are fixed from the following set of ‘default’ values: *T_i_* = 0, *α* = ∞, *μ* = *ν, E_B_* = ∞(*δ_n_* = 0). Each scenario can be used to elude kinetic traps and achieve a high assembly yield but how much time do these different strategies require?

In artificial self-assembly systems, the temporal supply of components can usually be controlled externally. This offers effective ways of regulating the assembly dynamics. To examine the potential of such supply-control strategies, we study two diametrically opposed cases. In the first case, all building blocks are supplied (activated) uniformly over a fixed time interval *τ* = 1/*α* at a constant influx rate *Nα*. By controlling *α* one can regulate the concentrations of monomers and hence the effective dimerization rate. In the second case, the different species are added in a defined temporal sequence (Fig. 1), which allows to favour specific assembly pathways by altering the order of the time points *T_i_* at which a species *i* is added (supply order).

Besides the binding rate *ν* that fixes the timescale, we are left with four control parameters, *E_B_*, *μ*, *α*, {*T_i_*}, which define different assembly scenarios (Fig. 1). In the reversible binding scenario, kinetic traps are avoided by ‘designing’ monomers with an optimal binding energy *E_B_* and resulting detachment rates *δ_n_*. This strategy is considered as the state of the art in DNA-brick-based self-assembly [11, 17, 34, 35]. In the dimerization scenario, the assembly process is controlled by the dimerization rate *μ*. A nucleation barrier *μ/ν* < 1 can be implemented for example by allosteric effects or with the help of enzymes (assembly factors) and is known to play a central role in many instances of biological self-assembly [19, 21, 38]. In the activation scenario, the assembly efficiency is controlled by an overall influx rate *α* without discrimination between species. Such a control of the availability of active monomers has been suggested as a means to effectuate the self-assembly of virus capsids [19], as well as other cellular macromolecular structures [20]. Finally, in the just-in-sequence (JIS) scenario, the monomers are supplied just-in-sequence with a favourably chosen assembly path by appropriate design of the supply order {*T_i_*}.

It should be noted that, although we consider a fully heterogeneous model system here, the dimerization, activation and reversible binding scenarios, which treat all species equivalently, are actually independent of the heterogeneity of the structure (if the particle number *N* is large enough, see SI). Only the JIS scenario requires at least two distinct species (see Methods). Therefore, the results of our time complexity analysis hold for general self-assembly systems independent of their heterogeneity.

## 3 Time complexity analysis

For each of the four scenarios, we investigated the minimal time required to achieve a target yield of 90% (denoted as 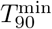). This requires to identify the optimal value of the respective control parameter that maximizes the time efficiency. The minimal assembly time is measured in units of the reactive timescale (*Cν*)^−1^, which defines the basic timescale of the process, and is inversely proportional to the concentration *C* = *N/V* of monomers per species in the reaction volume *V*. We are interested in the asymptotic dependence of 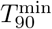 on the structure size *S* for *S, N* ≫ 1. In particular, while we have shown previously [12] that for small copy numbers *N*, the activation scenario is strongly influenced by stochastic effects, we here assume *N* to be large enough so that stochastic effects can be considered irrelevant. In this deterministic regime, we find that both the optimal control parameter and the minimal assembly time exhibit power-law dependencies on the size *S* of the target structure (Fig. 2). The corresponding exponents are referred to as the control parameter exponent *ϕ* and the (time) complexity exponent *θ*, respectively. Both exponents are scenario-specific and, moreover, depend on the dimensionality of the assembled structure, as discussed next for each scenario.

**Figure 2.**
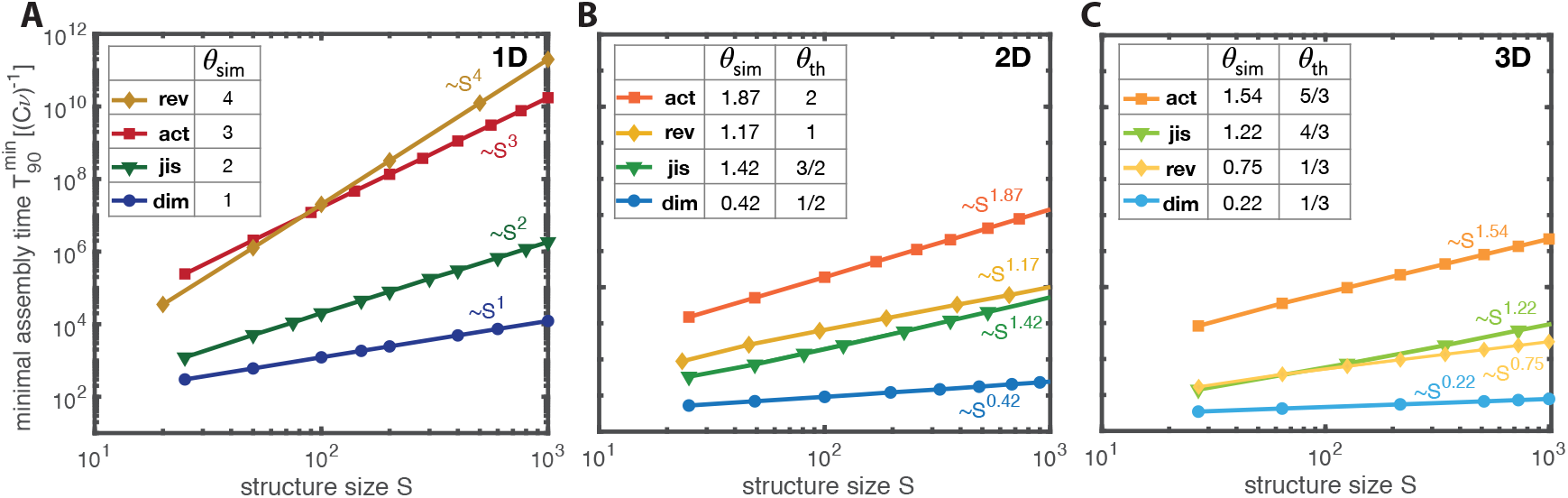
Time complexity. The minimal assembly time 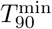 in the four scenarios in dependence of the size *S* of the target structure for different dimensionalities of the structure: (A), 1D; (B), 2D; (C), 3D. The minimal assembly time is measured in units of the reactive time scale (*Cν*)^−1^. Each data point represents an average over several independent realizations for the same parameter value (see Methods). We find power-law dependencies of the minimal assembly time on the size of the target structure. The corresponding time complexity exponents *θ*_sim_ resulting from the simulations are summarized in the table in each subpanel together with their theoretic estimates *θ*_th_ (see main text and SI). We indicate the scenarios as rev:=reversible binding, act:=activation, jis:=just-in-sequence and dim:=dimerization.

### 3.1 Reversible binding scenario

In the reversible binding scenario, the time complexity strongly depends on the dimensionality of the structure. For one-dimensional structures, the rate of monomer detachment is the same for all unfinished structures. Hence, it is not possible to selectively disfavour the growth of small structures by varying the binding energy *E_B_*. Instead, as we show in the SI, the effective time evolution of cluster sizes self-organizes into a diffusion process with an effective diffusion constant 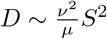. Hence, the assembly time for one-dimensional structures scales like the diffusive timescale 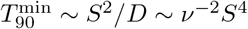 with time complexity exponent *θ* = 4, which agrees very well with the results obtained from stochastic simulations (Fig. 2A). In higher dimensions, large clusters are typically bound more tightly and hence become energetically favoured over clusters of small size, as illustrated in Fig. 3A. This creates an effective nucleation barrier, which allows one to strongly enhance the time efficiency compared to the one-dimensional case. However, in order to guarantee both high resource efficiency (high yield) and time efficiency, the binding energy must be fine-tuned to within few per cent of its optimal value (see Fig. 3B). Larger binding energies imply a lower nucleation barrier and lead to kinetic trapping, whereas lower binding energies progressively reduce the effective nucleation rate. By fine-tuning of the binding energy *E_B_*, we obtain the time complexity exponents *θ*_2*D*_ ∼ 1.19 and *θ*_3*D*_ ∼ 0.75 respectively, for the two- and three-dimensional cases (Fig. 2B,C). Note that the binding energy *E_B_* is measured in units of *k_B_T* and the detachment rate relative to the reaction rate (*Cν*) (see Fig. 3B and SI). Hence, the most feasible way to fine-tune the control parameter in experiments will be to adapt either the temperature or the monomer concentration.

### 3.2 Dimerization scenario

We then analysed the remaining irreversible assembly scenarios, setting *δ_n_* = 0. In the dimerization scenario, decreasing the dimerization rate *μ* disfavours initiation of new structures relative to the growth of existing structures. Figure 3C shows the corresponding transition from zero to perfect final yield, with *μ*_90_ indicating the rate at which a final yield of 90% is achieved. We find that the optimal rate *μ*_opt_ that minimizes the time required to achieve 90% yield is only slightly lower than *μ*_90_ and, for linear structures, scales as *μ*_opt_ ~ *νS^−^*^2^ (see inset, Fig. 3C). This result obtained from simulations is supported by scaling arguments; see SI. Since dimerization is the rate-limiting step, we expect that the assembly time will predominantly be determined by the total dimerization rate 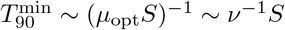. This estimate correctly predicts the time complexity exponent *θ* = 1 for linear structures (Fig. 2A).

Interestingly, in the dimerization scenario, the exponents for target structures of higher dimension can be related quite well to the one-dimensional case by a simple scaling argument: The number of possible binding partners of a globular structure with *s* particles is proportional to its surface area, and thus scales approximately as *s*^(*d−*1)/*d*^) where *d* is the dimensionality of the structure. Thus, defining an effective average growth rate *ν_S_ ~ ν S*(^*d−*1)/*d*^ for a target structure size *S* allows one to map higher-dimensional growth processes to an effective one-dimensional process along the radial coordinate. Replacing *ν* → *ν_S_* therefore translates the scaling laws for linear objects into approximate scaling laws for higher dimensional structures. This scaling idea for the dimerization scenario accurately yields the control parameter exponents for higher dimensional structures (see table in Fig. 3C) and only slightly overestimates the time complexity exponents in higher dimensions (Fig. 2B,C). These deviations may be attributed to the subleading contribution of the growth process to the total assembly time, which becomes more pronounced in higher dimensions.

**Figure 3.**
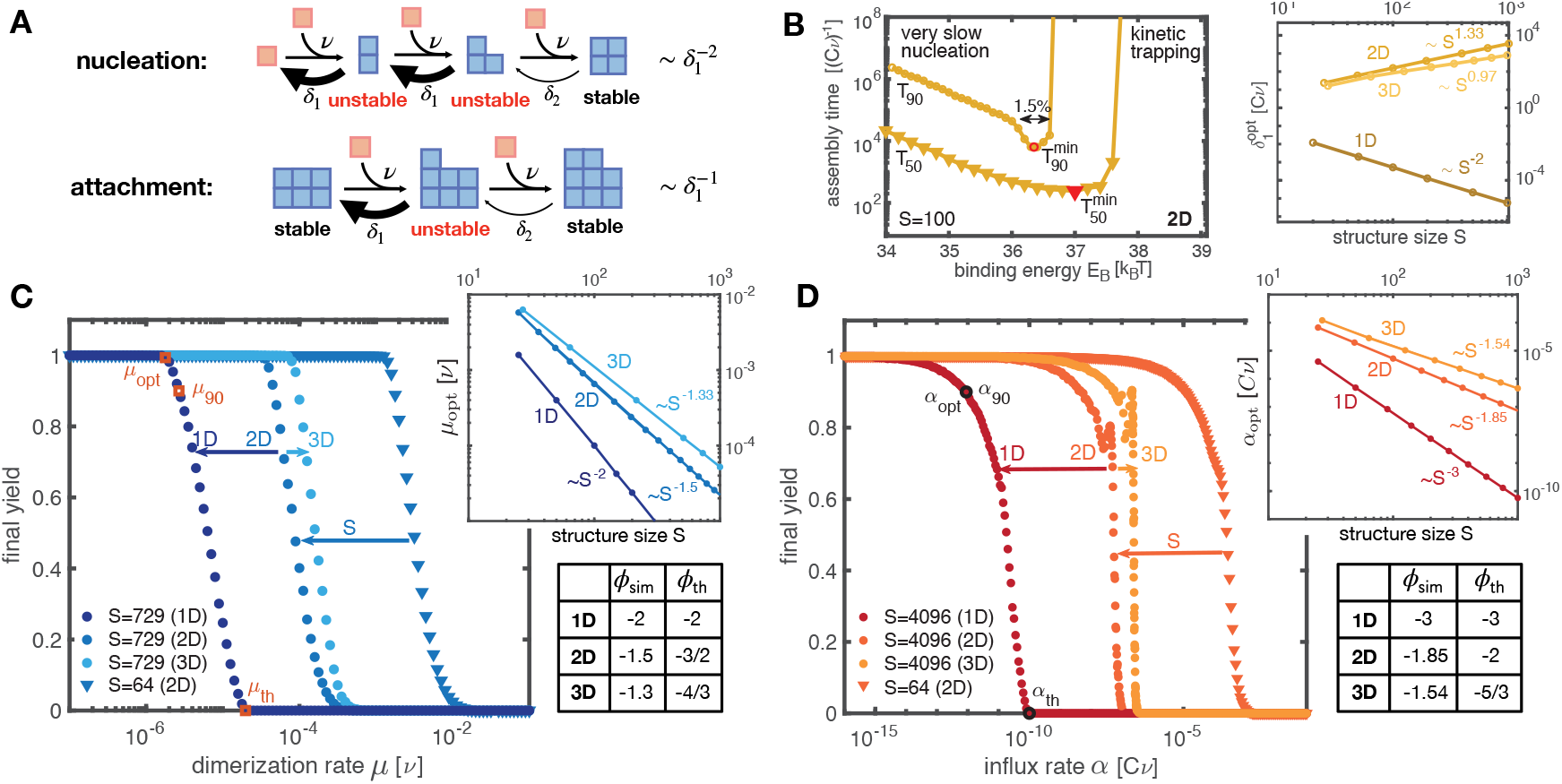
Reversible binding, dimerization and activation scenario. **A**, In the reversible binding scenario, (if *δ*_2_ ≪ *δ*_1_) the cluster evolution typically proceeds via stable intermediate states (in which all constituents form two or more bonds), whereas unstable intermediates are short-lived. Hence, nucleation is disfavoured relative to growth because nucleation proceeds via two unstable intermediate states whereas attachment proceeds only via one. **B**, Assembly time to achieve 50% yield (*T*_50_) and 90% yield (*T*_90_) plotted against the binding energy *E_B_* for two-dimensional target structures of size *S* = 100 (with preexponential factor *A* = 10^18^*Cν*). In order to achieve high yield with maximal time efficiency, *E_B_* must be fine-tuned to a narrow range (here ≈ 1, 4%) around its optimal value. Inset: The optimal detachment rate 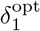 exhibits a power-law dependence on the structure size with exponent characterized by the dimensionality of the structure. **C**, **D**, Dimerization and activation scenario. The final yield in dependence of the dimerization rate (C) and the activation rate (D) for different sizes (symbols) and dimensionality (color shading) of the target structure. Data points represent averages over at least 20 independent realizations. Upon decreasing either the dimerization or activation rate, perfect final yield is achieved. For the leftmost transition we indicate the optimal parameter value *μ*_opt_ or *α*_opt_ that minimizes the time to achieve a yield of 90%. Insets show the dependence of the optimal parameter value on the structure size for different dimensionality. The corresponding control parameter exponents *ϕ*_sim_ are summarized in the table together with their theoretic estimates *ϕ*_th_ (see main text).

### 3.3 Activation scenario

In the activation scenario, nucleation is inhibited by controlling the concentration of available (or active) monomers. Decreasing the influx rate *α* reduces the momentary concentration of active monomers, and therefore reduces the effective dimerization rate. As in the dimerization scenario, this leads to a transition from zero to perfect final yield (Fig. 3D). However, the transition is not strictly monotonic but exhibits some small-scale peaks, whose origin is not entirely clear. From the stochastic simulations and scaling analysis (see SI) we infer for linear structures an optimal influx rate scaling as *α*_opt_ ~ *ν*^2^*S^−^*^3^ (inset, Fig. 3D). The control parameter exponents *ϕ* for higher dimensional structures derived from our rescaling argument are slightly larger than those obtained from simulations (table in Fig. 3D). As the monomers are activated over a time span 1/*α*, the time complexity exponents are the reciprocals of the parameter exponents (Fig. 2).

### 3.4 Just-in-sequence scenario

In the irreversible assembly scenarios discussed so far, all species are made available simultaneously. Consequently, excess nucleation of structures can only be suppressed by using a low dimerization or activation rate. In contrast, the JIS scenario favours specific assembly paths by regulating the order in which building blocks are supplied. The species supplied first in this temporal sequence define the nuclei for subsequent growth. Formation of other competing nuclei (dimers) during the assembly process is suppressed by the sequential delivery of building blocks, which ensures that mutual binding partners are supplied successively. Binding of newly added monomers to existing structures is therefore more likely than formation of new dimers. The frequency of competing nucleation events can be controlled by adjusting the interval Δ*T* between the equidistant time points *T_i_* at which subsequent ‘batches’ of monomers are provided. Longer time intervals increase the yield at the cost of a lower time efficiency. In order to minimize the total number of batches, we chose an ‘onion’ supply protocol, which allows structures to grow radially from the inside out, like the skins of an onion; see Fig. 4C. Furthermore, the time efficiency can be enhanced by using increasing, non-stoichiometric concentrations for the monomers in successive batches (Fig. 4B). Non-stoichiometric concentrations in a properly chosen ratio (see Methods and SI) reduce competition for resources between growing structures (Fig. 4A) and thereby greatly enhance the time efficiency, as well as robustness to extrinsic noise in the particle numbers supplied, especially for higher dimensional structures (cf. Fig. 4D,E). Therefore, non-stoichiometric concentrations are the key to successful implementation of the JIS strategy for higher dimensional structures. Since we assume equidistant time intervals Δ*T* between subsequent batches, the total assembly time is the product of Δ*T* (~*S*) and the total number of batches (~ *L* ~ *S*^1/*d*^), yielding the complexity exponents *θ* = 1 + 1/*d*, as shown in Fig. 2. In order to demonstrate the broad experimental applicability of the JIS supply strategy with a concrete example, we discuss in detail in the SI how the JIS strategy could efficiently be used to assemble artificial T=1 capsids.

**Figure 4.**
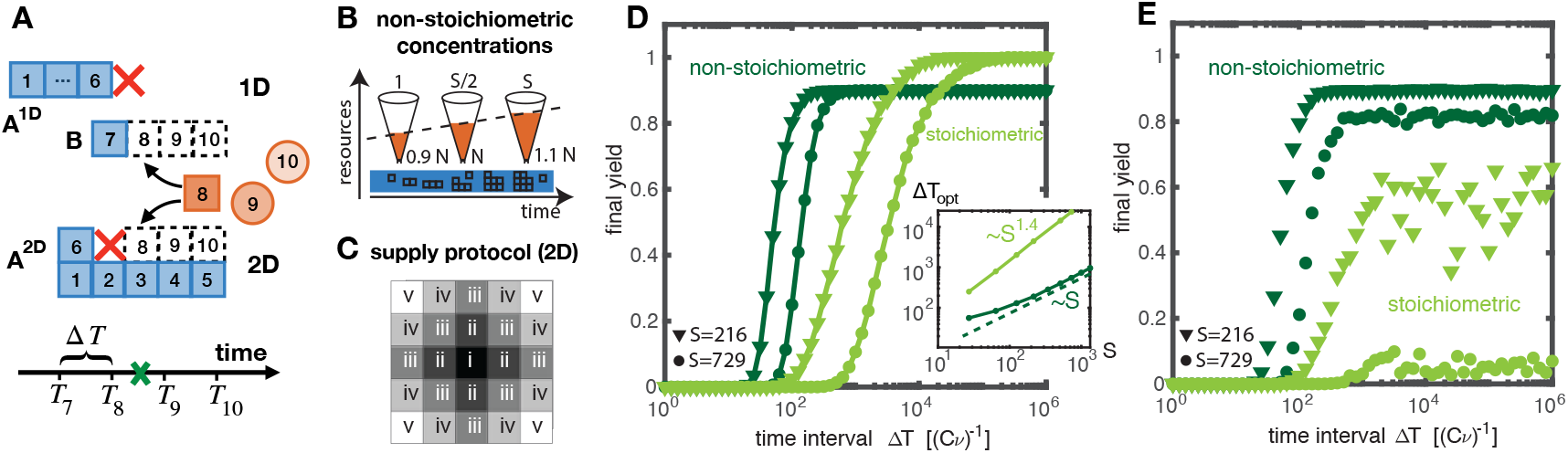
JIS scenario. **A**, In the JIS scenario, the different species are added sequentially; here, for illustration, in a linear sequence (*T*_1_ < *T*_2_ < *T*_3_ < …). Along the regular assembly paths, *A*_1*D*_ (1D) or *A*_2*D*_ (2D), additional dimers *B* can form competing for resources with the regular structures and thereby disrupting their growth. While for one-dimensional structures a disruption event prevents a structure *A*_1*D*_ from further growth, in higher dimensions both defective structures *A*_2*D*_ and *B* continue to grow thereby increasing competition for resources. **B**, Competition for resources can be alleviated by enhancing the amount of resources with each assembly step (non-stoichiometric concentrations, see Methods). For example, providing the first species in concentration 0.9*N* and increasing linearly up to 1.1*N* for the last species strongly enhances assembly efficiency (D) and robustness (E). **C**, Parallel supply protocol illustrated for a 2D structure of size *S* = 25 causing the structures to grow radially in an ‘onion-skin’ like fashion. Roman numbers indicate the order in which species are supplied. Species with identical numbers (‘onion skins’) are supplied simultaneously in ‘batches’. **D**, When using non-stoichiometric concentrations (see (B) and Methods), high yield can be achieved with a shorter time span Δ*T* between subsequent batches, exhibiting a smaller control parameter exponent (inset) as compared to the case of stoichiometric concentrations. Simulations were performed for 3D structures with *N* = 10^4^ − 10^5^. **E**, External noise in the concentrations jeopardizes the yield when stoichiometric concentrations are used, whereas non-stoichiometric concentrations are much more robust. Here, for each species we assumed a coefficient of variation CV = 0.1% with average particle numbers as in (D).

## 4 Discussion

Figure 2 shows the dependence of the minimal assembly time on target structure size, together with the resulting time complexity exponents for the different scenarios and dimensionalities. All exponents decrease with increasing dimensionality of the target structure and can even change their relative order. Remarkably, the exponents are robust to various modifications of the model such as heterogeneous binding rates, modified boundary conditions or altered definitions of the assembly time (see SI). Similarly, advanced protocols like annealing or different forms of monomer input in the activation scenario leave the exponents invariant (see SI). This invariance shows that the time complexity analysis yields a reliable and robust characterization of self-assembly processes. Furthermore, the invariance of the parameter exponents allows for an optimal control strategy to be identified in dependence of the size of the target structure in each of the four scenarios.

The dimerization scenario turns out to be the most time-efficient scenario in all dimensions. Controlling the dimerization rate is efficient as it allows to initiate just as many structures as are needed, followed by a rapid growth phase if all particles are readily available. For linear structures, the supply-control strategies rank second and third, with coordinated supply in the JIS scenario being more efficient than uncoordinated supply in the activation scenario. Reversible binding is the least efficient approach to assembling large linear structures, but it is efficient for the assembly of higher dimensional structures and then becomes competitive with the JIS scenario, slightly outperforming it for large structure sizes.

In conclusion, our time-complexity analysis of self-assembly describes lower bounds for the required assembly time as a function of the target structure size. Furthermore, it accomplishes a robust description of how the parameters of the system must be controlled in order to achieve optimal time and resource efficiency. The analysis enables us to compare the efficiency of different assembly scenarios. In computer science, the complexity of a computational problem is defined as the complexity of the fastest algorithm available to solve it [28]. Among the assembly scenarios discussed here, limiting the dimerization rate defines the fastest assembly process, and might thereby determine the time complexity of self-assembly (of course, we cannot exclude the possibility of even faster assembly strategies). Experimentally, however, controlling the dimerization rate is difficult, as it effectively requires building blocks that exhibit allosteric binding effects. So far, experiments have typically resorted to rendering binding reactions reversible [17, 25,33–36]. Our analysis shows that this common approach is time efficient for the assembly of higher dimensional structures. However, to be truly competitive, fairly precise tuning of bond strengths, temperature and the concentration is required (see SI). Our analysis suggests that a supply control strategy like the JIS scenario is a promising alternative that offers similar or better time efficiency using irreversible self-assembly. As a significant advantage, this strategy does not rely on sophisticated properties of the building blocks (like allosteric effects or fine-tuned bond strengths) but only on temporal supply control and hence on parameters that might be more amenable to regulation and adaptation in experiments. Compared to the current state-of-the-art approach via reversible reactions, these irreversible assembly schemes might thus provide a complementary and more versatile strategy for assembling complex structures, requiring control over relative concentrations, rather than fine-tuning of the molecular details. Importantly, the idea underlying the JIS scenario entails a rather specific design principle for efficient irreversible assembly protocols of complex nanostructures (‘batches without mutual binding partners’); We demonstrate in the SI how this principle is applied exemplarily for the assembly of artificial T=1 capsids. This design principle thereby provides a clear path towards the experimental realization of the JIS scenario, suggesting that the strategy will be broadly applicable to the assembly of artificial structures.

An interesting question for future research concerns the prospects for spatiotemporal supply control, i.e. controlling not only the time interval but also the site at which monomers are injected into a spatial system, for further enhancement of the time efficiency. Moreover, it would be interesting to consider the time complexity of assembly schemes like hierarchical self-assembly [2, 16, 23, 31], which include polymer-polymer interactions. Finally, other potentially important aspects of self-assembly include susceptibility to errors in the case of reduced binding specificities or defective particles, as well as robustness to stochastic effects for small copy numbers. It might therefore be instructive to test the different assembly scenarios with respect to these factors too.

## 5 Methods and Materials

### 5.1 Simulation

Particle-based, stochastic simulations of the reaction kinetics of the system were performed using Gillespie’s algorithm [13]. Our C++ code will be made available upon request. In the following we discuss the parameter settings and some particularities of the individual scenarios that are relevant for their simulation. In the Supplementary text below, we also discuss the moment equations that result from the stochastic Master equation of the system and the “method of homogenization”. This method exploits the equivalence of the system to a homogeneous system with indistinguishable species, requiring a lower total number of particles and thereby increasing the efficiency of the simulation (in particular that of the activation scenario).

### 5.2 Reversible Binding Scenario

For the reversible binding scenario, the parameters were set as follows: *μ* = *ν* = 1, *α* = ∞ (i.e. all monomers are available right from the outset), *T_i_* = 0 ∀*i* and a variable binding energy per contact *E_B_* that fixes the detachment rates according to 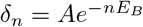 (Arrhenius’ law). We fixed the pre-exponential factor at *A* = 10^18^ *Cν*, which appears to be a realistic choice in the light of typical experimentally measured values for A [4, 26]. However, we confirmed that the choice of the constant *A* does not qualitatively affect our results (in particular it does not affect the exponents) as long as *A* is large, and hence *δ*_1_ ≫ *δ_n>_*_1_. If *A* is small (for example *A* = 10^6^*Cν* or smaller), or when *δ_n_* values are chosen independently of one another, the minimal assembly time and the measured exponents can differ slightly, as *δ*_2_ is no longer negligible compared to *δ*_1_ (see Fig. S1).

We simulated the reversible binding scenario with particle number *N* = 500. It is important that *N* is chosen large enough, because for small *N* the measured assembly time fluctuates very strongly between independent runs and the average assembly time increases with *N*. Only if *N* is large enough does the average assembly time (measured relative to the reactive timescale *Cν* as in Fig. 2) converge and become independent of *N*. We verified that for *N* = 500 the remaining *N*-dependence is negligible. Alternatively, the method of homogenization described in the SI can be used to reduce the role of fluctuations resulting from finite particle numbers and therefore allows the system to be simulated with fewer particles. In particular, the reversible binding scenario in one dimension can be simulated faster and more accurately in this way with a fivefold lower total particle number (*N*_tot_ = 100*S*).

Generally, simulation of the reversible binding scenario is computationally much more expensive than that of the irreversible scenarios, since many more steps are generally needed owing to the fast detachment processes. Partly, a single run needed several billion Gillespie steps to complete. It is therefore useful to reduce the particle number in the simulations, as long as the results remain accurate.

### 5.3 Dimerization Scenario

For the dimerization scenario we used *α* = ∞, *T_i_* = 0 ∀*i, δ_n_* = 0 and a variable dimerization rate *μ* as well as *N* = 1000. The dimerization scenario can be simulated most efficiently, because far fewer steps are needed due to the irreversibility of binding reactions. Furthermore, stochastic effects do not play an important role [12], so *N* can be chosen to be relatively small. Conversely, fluctuations in the assembly time between independent runs decrease with increasing *N*, allowing for greater accuracy in the determination of the exponents.

### 5.4 Activation Scenario

We defined the activation scenario by *μ* = *ν* = 1, *T_i_* = 0 ∀*i, δ_n_* = 0 and a variable influx rate *α*. Since the momentary concentration of active monomers is generally small for a low influx rate, the activation scenario is strongly affected by stochastic effects (see reference [12] for details). Furthermore, the magnitude of these stochastic effects strongly depends on the number of species, and hence on the size *S* of the target structure. Consequently, depending on *S*, a large number of particles *N* may be required to achieve a yield ≥ 90% in the activation scenario. By “homogenizing” the system, i.e. treating species as indistinguishable and simulating a homogeneous system instead of a heterogeneous system as described in the SI, the computational cost of the simulation can be drastically reduced using a much smaller total particle number. In the case of periodic boundary conditions of the structure, the homogenized simulation can be shown to be exact, in the sense that it reproduces the yield and assembly time obtained for the heterogeneous system in the limit of large *N* (see supplementary text). In the case of open boundaries of the structures (as assumed in the main text), “homogenization” yields a very accurate approximation (see Fig. S2). We exploited this method to simulate the activation scenario efficiently and accurately with a total number of particles *N*_tot_ = 1000*S*, as in the dimerization scenario.

### 5.5 Just-in-Sequence Scenario

For the JIS scenario, we set *μ* = *ν* = 1, *α* = ∞, *δ_n_* = 0 and control the time points *T_i_* at which the different species are supplied. Species with identical *T_i_* define a ‘batch’. We only considered the case of equidistant intervals Δ*T* between successive batches. The supply protocol (see Fig. 4C) assigns the species to the batches and specifies the concentrations in which the species are supplied. In this work, we exclusively used the “onion-skin supply protocol” depicted in Fig. 4C, where structures grow radially from the center outwards. This protocol minimizes the total number of batches. As discussed in the main text, in the JIS scenario, choosing the concentrations of the species in specific, non-stoichiometric ratios is crucial in reducing competition for resources among the growing structures and enhancing the efficiency of assembly. In order to compensate for the increasing number of clusters that form through excess dimerization events, the number of resources supplied is increased with each batch. This comes at the price, insofar as the maximum yield is limited to a value less than 1 corresponding to the number of initial seeds. The desired effect is that each species can be provided in an amount that allows all the structures currently present in the system to grow, thus reducing competition for resources to a minimum. The optimal concentration of a species is therefore determined by the total number of structures formed during previous assembly steps that are capable of binding the species that has just been supplied. More precisely, for each species provided in the *b*^th^ batch, we supply a number

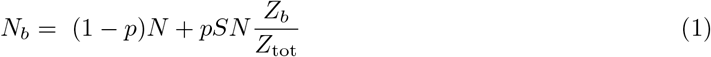

of monomers, where *Z*_1_ = 0 and *Z_i_* < *Z_j_* for *i* < *j*, see below. The first contribution, (1 – *p*)*N*, which is identical for all species, is the basal particle number, which defines the maximum number of complete structures that can be built. The second contribution is the excess concentration, which provides additional resources for the growing total number ~ *Z_b_* of complexes that have already formed through excess dimerization events. Here, pSN with *p* < 1 is the total amount of resources that is distributed unevenly among the species, and *Z_b_/Z*_tot_ is the fraction of that amount assigned to the individual species supplied in the *b*^th^ batch. The normalization factor 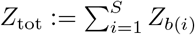, with *b*(*i*) denoting the batch number of species *i*, sums the *Z_b_* over all species, and thereby fixes the average particle number 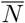 per species: 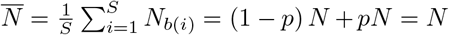. The basal fraction of resources (1 − *p*) determines the maximum yield, and hence should be at least 0.9 to meet our criterion for the assembly time. We found that *p* = 0.07 minimizes the assembly time *T*_90_ and, therefore, we used this value in the simulations.

The success of the JIS strategy crucially depends on the choice of the numbers *Z_b_*. Optimally, the numbers *Z_b_* should reflect the number of the excess complexes relevant for a species supplied in the *b*^th^ batch (see Fig. S3). Approximately, the number of previous excess dimerization events will be proportional to the total number of species supplied previous to the *b*^th^ batch, i.e. provided by the batches 1 to *b* − 1. Since in the onion-skin protocol, species with batch number less than *b* form a *d*-dimensional volume (see Fig. 4C), for large *b* we obtain approximately: *Z_b_* ~ *b^d^*. Correcting this count for small *b* (see Fig. S3) we can further improve the efficiency by setting:

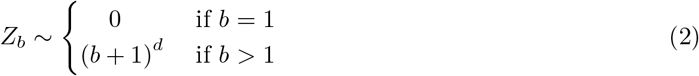

for two- and three-dimensional structures and *Z_b_* ∼ (*b* − 1) in the 1D case. It might be possible to improve the efficiency further by assigning particle numbers *N_s_* individually for each species, rather than identically for all species in the same batch. However, we already achieve very good results with this choice of *Z_b_*. On the other hand, with all species in a batch being supplied in identical particle numbers, those species could likewise be indistinguishable. In this way, a regular target structure could be designed with only two distinct species, which alternately assemble the “skins of the onion” (the “homogenized” version of the JIS scenario; also see the example on capsid assembly in SI section 4). Furthermore, note that, if the particle numbers *N_b_* for the different species are chosen appropriately on average, the system becomes robust to external noise up to a certain limit (see Fig. 4E and Fig. S7B).

For reasons of computational efficiency we would like to simulate the system with a small (average) particle number *N*. Note, however, that the implementation of non-stoichiometric concentrations requires a minimum *N* due to the discreteness of particle numbers: In order to ensure that the right-hand side of Eq.(1) reasonably maps onto integer values for the numbers *N_b_*, the factor *pSN/Z*_tot_ that multiplies *Z_b_* should be of the order at least 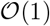. In order to find a rough condition for *N*, we therefore estimate the normalization factor *Z*_tot_:

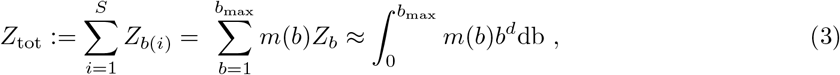

where in the second step we change the sum over *species* to a sum over *batches*, with *m*(*b*) denoting the number of species in the *b*^th^ batch (‘density of species’) and 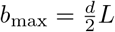 being the total number of batches (see Fig. 4C). Note that, in the onion-skin protocol, species with the same batch number lie on rhomboidal shapes around the center species. Furthermore, the densities are symmetric about b_max_/2 (batches ii and iii have the same densities as v and vi, respectively, in the supply protocol depicted in Fig. 4C). Hence, we approximate the density of species by

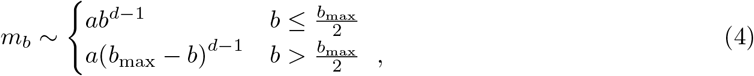

where the constant *a* is determined from the condition 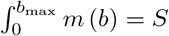. Performing the calculation yields *Z*_tot_ ~ *S*^2^. Hence, in order to guarantee that the prescribed ratios of the particle numbers *N_b_* can be met, the average particle number should be 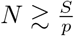.

We used *N* = 10^4^ in our simulations of the JIS scenario with non-stoichiometric concentrations, with *p* = 0.07 and a structure size *S* of maximally 10^3^. By simulating individual runs with a larger particle number *N* = 10^5^, we verified that the *N*-dependence of the assembly time is negligible for *N* ≥ 10^4^. The simulations of the JIS scenario with stoichiometric concentrations were performed with *N* = 10^5^, because the larger time intervals Δ*T* led to very small momentary concentrations, and hence required a larger overall particle number to achieve *N*-independent assembly times.

### 5.6 Determination of 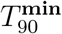 and the optimal parameter

In order to determine the minimal assembly time for a specified scenario and target structure, we first varied the respective control parameter roughly to find an estimate for its optimal value that minimizes the assembly time in the simulation. Afterwards, we sampled the parameter range around the estimated parameter value thoroughly by varying the control parameter in equidistant increments of approximately 2-4 percent precision. For each parameter value, the assembly time was averaged over several independent runs (50-100 for the irreversible scenarios and 5-50 for the reversible binding scenario). The minimal assembly time 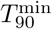 was then determined as the minimum of the averaged assembly times, and the corresponding parameter value was chosen as the optimal parameter value. If the minimum of the assembly times was attained at the boundary of the sampled parameter range, we increased the range in the direction of the respective boundary and simulated additional parameter values. We repeatedly increased the range (or modified the parameter estimate) until we found a minimum that was attained somewhere in the middle of the sampled range to ensure that the global minimum has been identified.

## 6 Acknowledgements

We thank Patrick Wilke and Philipp Geiger for stimulating discussions. This research was supported by the German Excellence Initiative via the program ‘NanoSystems Initiative Munich’(NIM) and was funded by the Deutsche Forschungsgemeinschaft (DFG, German Research Foundation) under Germany’s Excellence Strategy – EXC-2094–390783311 as well as under – Project-ID 364653263 – TRR 235. FMG and IRG were supported by a DFG fellowship through the Graduate School of Quantitative Biosciences Munich (QBM).

## Notes

### Competing Interest Statement

The authors have declared no competing interest.

