## Supplementary Information for "The Time Complexity of Self-Assembly"

**CONTENTS**

A. Supplementary Information 3

1. Moment equations and the method of 'homogenization' 3
   The method of 'homogenization' 4

2. Scaling theory 7
   Reversible binding for 1D structures 7
   Irreversible scenarios and reversible binding for 2D/3D structures 11
    Activation and dimerization scenario 12
    JIS scenario 13
    Reversible binding for 2D and 3D structures 15

3. Robustness to model modifications 17
   Structures with periodic boundaries 17
   Heterogeneous binding rates 17
   Reduced resource efficiency 18
   Annealing (reversible binding scenario) 18
   Alternate input functions (activation scenario) 19

4. Experimental JIS supply protocol for the assembly of an artificial T=1 capsid 21

B. Supplementary Figures 24

References 31

### A. Supplementary Information

In this Supplementary Information (SI), we first discuss the moment equations resulting from the stochastic Master equation describing the kinetics of the self-assembly system. With the help of the moment equations we motivate the 'method of homogenization', a powerful approximation that was used to simulate the activation scenario efficiently. Subsequently, we derive analytic estimates for the time complexity and control parameter exponents using mathematical calculations and scaling arguments. These analytic estimates for the exponents are the basis for the 'theoretical values' presented in the main text. Afterwards, we demonstrate that our results, in particular the time complexity and control parameter exponents, are robust to modifications of the model and variations in the parameters. Finally, in order to demonstrate the broad applicability of the just-in-sequence scenario, we show how the supply strategy can be used in practice for the concrete example of artificial T=1 capsid assembly.

#### 1. MOMENT EQUATIONS AND THE METHOD OF 'HOMOGENIZATION'

Here we show the moment equations resulting from the stochastic Master equation that describe the assembly kinetics for one-dimensional structures. The higher dimensional cases are conceptually similar to the one-dimensional case but do not allow for a simple representation of all possible cluster configurations. Therefore, we restrict ourselves to illustrating the mathematical framework only for the 1D case. The moment equations are subsequently used to show that for structures with periodic boundaries, the heterogeneity (distinguishability of species) is irrelevant in the limit of large $N$. This is the basis of our 'method of homogenization', which exploits the equivalence between heterogeneous and homogeneous systems in order to increase the efficiency of the simulations.

For one-dimensional structures, each possible kind of polymer can be characterized by two variables: the length $\ell$ of the polymer, and the monomer species $s$ at its right end which will be referred to as the species of the polymer. We denote by $n_\ell^s(t)$ with $2 \leq \ell < L$ and $1 \leq s \leq S$ the number of polymers of size $\ell$ and species $s$ in the system at time $t$. Furthermore, $n_0^s$ and $n_1^s$ denote the number of inactive (not yet added) and active monomers of species $s$, respectively, and $n_L$ the number of complete structures.
The subsequent set of equations can then be interpreted in two different ways: Either all terms with a species index (upper index) outside the range $1 \leq s \leq S$ are considered as zero or species indices are taken modulo S. The first case describes the self-assembly of structures with an open, non-periodic boundary (the case considered in the main text). In contrast, the second case describes the assembly process of a periodic structure, i.e. a ring in this 1D case. We show in section 3 of this SI that the choice of the boundary condition only has a small effect on the assembly time and, in particular, does not affect the control parameter and time complexity exponents. By $\langle ... \rangle$ we indicate (ensemble) averages. The

system governing the evolution of the first moments (the averages) of the $\{n_\ell^s\}$ is then given by:

$$\frac{d}{dt}\langle n_0^s\rangle = -\alpha\,\Theta(t-T_s)\Theta(T_s+1/\alpha-t)\,, \tag{1a}$$

$$\begin{aligned}\frac{d}{dt}\langle n_1^s\rangle &= \alpha\,\Theta(t-T_s)\,\Theta(T_s+1/\alpha-t) - \mu\left(\langle n_1^s n_1^{s+1}\rangle + \langle n_1^s n_1^{s-1}\rangle\right)\\ &\quad - \nu\sum_{\ell=2}^{L-1}\left(\langle n_1^s n_\ell^{s+\ell}\rangle + \langle n_1^s n_\ell^{s-1}\rangle\right) + \delta\sum_{\ell=2}^{L-1}\left(\langle n_\ell^{s+\ell-1}\rangle + \langle n_\ell^s\rangle\right)\,,\end{aligned} \tag{1b}$$

$$\frac{d}{dt}\langle n_2^s\rangle = \mu\,\langle n_1^{s-1}\,n_1^s\rangle - \nu\,\left(\langle n_1^{s-2}\,n_2^s\rangle + \langle n_2^s\,n_1^{s+1}\rangle\right) + \delta\,\left(\langle n_3^s\rangle + \langle n_3^{s+1}\rangle - 2\langle n_2^s\rangle\right)\,, \tag{1c}$$

$$\begin{aligned}\frac{d}{dt}\langle n_\ell^s\rangle &= \nu\,\left(\langle n_1^{s-\ell+1}\,n_{\ell-1}^s\rangle + \langle n_{\ell-1}^{s-1}\,n_1^s\rangle - n_1^{s-\ell}\,\langle n_\ell^s\rangle - \langle n_\ell^s\,n_1^{s+1}\rangle\right)\\ &\quad + \delta\,\left(\langle n_{\ell+1}^s\rangle + \langle n_{\ell+1}^{s+1}\rangle\right)\mathbf{1}_{\{\ell\le L-2\}} - 2\delta\langle n_\ell^s\rangle\,, \qquad 3\le\ell<L\,,\end{aligned} \tag{1d}$$

$$\frac{d}{dt}\langle n_L\rangle = \nu\,\sum_{s=1}^{L}\left[\langle n_1^{s-L+1}\,n_{L-1}^s\rangle + \langle n_{L-1}^{s-1}\,n_1^s\rangle\right]\,. \tag{1e}$$

Eq. (1a) and the first term in Eq. (1b) describe the influx of monomers of species $s$ into the system starting at time $T_s$ until $T_s+\frac{1}{\alpha}$. Here, $\Theta$ denotes the Heaviside function. Besides the influx of monomers, the temporal change in the number of active monomers (Eq. (1b)) is governed by the following processes: dimerization of monomers at rate $\mu$, binding of monomers to the left and to the right end of existing polymers at rate $\nu$ and detachment of monomers from the left and right end of polymers with rate $\delta$.

Equations (1c) and (1d) describe the dynamics of dimers and larger polymers of size $3\le\ell<L$, respectively. The terms account for dimerization of active monomers as well as all possible kinds of reactions of polymers with monomers, together with detachment of monomers from polymers. The indicator function $\mathbf{1}_{\{\ell\le L-2\}}$ in Eq. (1d) (which equals 1 if the condition $\ell\le L-2$ is satisfied and 0 otherwise) excludes source terms that would account for detachment from completed structures, which are assumed to be stable. Finally, the complete structures form an absorbing state and, therefore, include only the respective gain terms (cf. Eq (1e)).

**The method of 'homogenization'**

For sufficiently large particle numbers $N$, correlations between the particle numbers $\{n_\ell^s\}$ in Eq. (1) can be neglected and the two-point correlator can be approximated as the product of the corresponding mean values (mean-field approximation):

$$\langle n_i^s n_j^k\rangle = \langle n_i^s\rangle\langle n_j^k\rangle\ \forall s,k \tag{2}$$

Note that, in the case of periodic boundary conditions, all species have equivalent binding properties. Mathematically, this is reflected by the invariance of Eq. (1) with respect to relabelling the upper indices if $T_i = T_j \; \forall i, j$. This symmetry of the system allows us to drop the distinction by species and to define the homogeneous concentrations

$$\langle n_\ell^s \rangle = \langle n_\ell^k \rangle := c_\ell \, V \quad \forall s, k, \tag{3}$$

where $V$ is the reaction volume. In this way, Eq. (1) reduces to a system of rate equations for a homogeneous (one species) system in the deterministic limit $N \to \infty$. Hence, in the case of periodic boundary conditions, if the particle number $N$ is large, the heterogeneity (distinguishability of species) is irrelevant; also see reference [1] for more details. This holds true for the activation, dimerization and reversible binding scenario where $T_i = T_j$.
The equivalence of species no longer holds exactly in the absence of periodic boundary conditions because then the species at the boundary of the structure violate the symmetry. However, the symmetry still holds approximately and the heterogeneous system can well be approximated by a corresponding homogeneous system for large $N$. Figure S2 shows that this approximation is indeed very accurate by comparing the deterministic behavior for systems with small structure size $S$.
The advantage of a homogeneous over a heterogeneous system is that stochastic effects arising from fluctuations in the concentrations of the different species are suppressed, [1]. Hence, in order to observe deterministic behavior, a smaller total number of particles $SN$ is required for homogeneous systems, increasing the efficiency of simulations. We exploit this increase of efficiency in our simulations of the activation scenario, where stochastic effects are particularly strong [1]. Instead of simulating the heterogeneous system, we implemented an (approximate) 'homogenization' method where only a single species, which can occupy any site in the structure, is simulated. To retain the asymmetry in the structure geometry itself (e.g. between bulk and boundary sites), the positions in the structure are still distinguished (e.g. there are boundary sites with fewer neighbors). In this sense, 'homogenization' can be thought of as simulating the dynamics of an 'average' species with reduced fluctuations. In the simplest way, a heterogeneous simulation can be 'homogenized' in two easy steps:

1. Make monomer creation and annihilation act on all species simultaneously (i.e. if an (unbound) monomer of one species is added or subtracted, add or subtract one for all other species as well),

2. rescale the influx rate $\alpha$ and dimerization rate $\mu$ by $S^{-1}$.

The first step constrains all species to equal concentrations while the second rescales the rates as if there were only a single species. Computationally, however, it is more efficient to simulate only one monomer species explicitly instead of acting on $S$ species simultaneously.

Note that, in principle, one could also solve the corresponding mean-field equations in order to investigate the system's deterministic behavior for large $N$. This approach, based on a description via ordinary differential equations, however, requires a characterization of all possible configurations of the clusters ('state-based' approach), which is not feasible for the higher dimensional systems due to the huge number of possible configurations. In contrast, 'homogenization' allows to stick with a particle-based description and hence avoids the necessity of characterizing all possible cluster configurations.

### 2. SCALING THEORY

In this section, we provide a mathematical scaling analysis in order to derive the characteristic exponents for the four scenarios analytically, supporting our numerical findings. We first discuss the reversible binding scenario for one-dimensional structures, followed by a unified approach to the irreversible scenarios as well as the reversible binding scenario for higher dimensional structures. Note that only the one-dimensional reversible binding scenario is fully reversible, while in the higher dimensional cases one can identify quasi-stable intermediate assembly products that form irreversibly. Exploiting the (stepwise) irreversibility of the assembly kinetics allows to analyze reversible binding for higher dimensional structures together with the irreversible scenarios in a unified approach, whereas reversible binding in one dimension needs to be analyzed separately.

#### Reversible binding for 1D structures

To mathematically analyze the scaling behavior of the one-dimensional reversible binding scenario, we need to identify the optimal value of the detachment rate $\delta := \delta_1$ that minimizes the time taken to achieve a yield of 90%, depending on the size of the target structure. Since generally several unfinished structures exist at the same time and thereby compete for resources when growing, an exact analysis requires knowledge of the full temporal evolution of the polymer size distribution, which is very hard to obtain. Therefore, we will make two simplifying assumptions to obtain the scaling behavior: First, we employ a quasi-stationarity assumption, $\partial_t m = 0$, for the monomer concentration. While this may seem to be a rather drastic postulate, the idea is rather intuitive: During the assembly process, structures grow by consumption of monomers and, vice versa, the number of monomers increases due to their detachment from structures. As a result, in the limit of large structure sizes where many attachment and detachment events occur before any structure is completed, the concentration of monomers adjusts itself over time in such a way that attachment and detachment roughly balance and the monomer concentration is constant. As we will show more explicitly below, in this case the polymer size distribution corresponds to a random walk on a one-dimensional lattice with constant hopping rates. To proceed, we then make a second, important assumption: We postulate that the scaling of the time to obtain a yield of 90% is the same as the scaling of the mean first-passage time of the approximate random walk to reach the absorbing boundary at $x = S$ (complete structure). This amounts to assuming that growth of structures is the time-limiting step and that the corresponding timescale does not change considerably over the course of the assembly process, e.g. the times to obtain 50 or 90% yield scale similarly with the structure size. With these assumptions, we identify the time complexity exponent to be 4 and the control parameter exponent to be -2, as we will outline in more detail in the following.

In the reversible binding scenario, we have $T_s = 0 \; \forall s$ and $\alpha \to \infty$. With the reaction rate $\nu$, the dimerization rate $\mu$ and the detachment rate $\delta$, the deterministic equations for the temporal evolution of the concentrations are (see Eqs. 1 for the general case):

$$\begin{aligned}
\partial_t m &= -2\mu m^2 - 2\nu m \sum_{j=2}^{S-1} c_j + 2\delta \sum_{j=2}^{S-2} c_j \\
\partial_t c_2 &= \mu m^2 - 2\nu m c_2 - \delta c_2 + 2\delta c_3 \\
\partial_t c_i &= 2\nu m(c_{i-1} - c_i) - 2\delta(c_i - c_{i+1}) \quad i = 3, \ldots, S-2 \\
\partial_t c_{S-1} &= 2\nu m(c_{S-2} - c_{S-1}) - 2\delta c_{S-1} \\
\partial_t c_S &= 2\nu m c_{S-1}
\end{aligned} \tag{4}$$

where $m$ is the number of monomers per species and $c_i$ the number of $i$-mers. Defining $K = \sum_{j=2}^{S-1} c_j$ to be the number of unfinished complexes, the temporal evolution for the monomers is given by

$$\partial_t m = -2\mu m^2 - 2(\nu m - \delta)K.$$

In the quasi-stationary limit, $\partial_t m = 0$, the evolution of the polymer-size distribution $\partial_t c_i$ can be identified with a random walk on a one-dimensional lattice with constant hopping rates $2\nu m$ to the right and $2\delta$ to the left, corresponding to monomer attachment and monomer detachment, respectively (see also the deterministic analogue in Eq. 4). Since completed structures are stable, the right end at $i = S$ is absorbing, implying that $c_S = 0$ or, in the continuum limit, $c(l = S) = 0$. Furthermore, we assume that all particles are provided at $t = 0$ at the left end $l = 0$[1]. The last two points imply that the polymer concentration $c(t, l)$ decreases over time. As a measure for the quasi-stationary properties of the system, we therefore consider the temporally integrated concentration $I(l) = \int_0^\infty \mathrm{d}t\ c(t, l)$.

In the continuum limit, Eq. 4 becomes $\partial_t c(t, l) = -2(\nu m - \delta)\partial_l c(t, l) + (\nu m + \delta)\partial_l^2 c(t, l)$. Using that $c(t \to \infty, l) = 0 \forall l$ and $c(0, l) = 0 \forall l > 0$, the integrated concentration satisfies $v\partial_l I(l) = D\partial_l^2 I(l)$ where

$$v = 2(\nu m - \delta)$$

is the drift coefficient and

$$D = \nu m + \delta$$

[1] Since we are interested in the limit of large $S$, we approximate $S - 2 \approx S$ and, thus, do not distinguish whether particles are injected at $l = 0$, $l = 1$ or $l = 2$.

is the diffusion constant of the random walk. Its solution is given by

$$I(l) = C(1 - e^{v(l-S)/D})$$

where $C$ is an integration constant that is related to the number of injected particles. It will, however, not be relevant for the calculation of the first-passage time.

We will use the integrated concentration to calculate the time-averaged mean size of unfinished polymers. This quantity is helpful to determine the number of monomers self-consistently as conservation of particles requires $m + \sum_{j=2}^{S} jc_j = N$. Before yield sets in this can be rewritten as $m + \sum_{j=2}^{S-1} jc_j = N$. Furthermore, the sum can be expressed in terms of the average polymer size of unfinished polymers $\langle j \rangle$ as $\sum_{j=2}^{S-1} jc_j = \langle j \rangle \sum_{j=2}^{S-1} c_j = \langle j \rangle K$. In the continuum limit, we find the following self-consistency equations:

$$N = m + \langle l \rangle K \tag{5}$$

$$\langle l \rangle = \frac{\int_0^S \mathrm{d}l \; lI(l)}{\int_0^S \mathrm{d}l \; I(l)} = -\frac{D}{v} + \frac{S^2 v}{2(Sv + D(-1 + e^{-Sv/D}))}. \tag{6}$$

From the quasi-stationarity condition $\partial_t m = 0$, we furthermore find

$$m^2 + \frac{\mu}{\mu} mK - \frac{\delta}{\mu} K = 0. \tag{7}$$

Taken together, we have three conditions 5, 6 and 7 to determine three unknown variables $m$, $K$ and $\langle l \rangle$ self-consistently (for fixed $\delta$). Furthermore, we have another unknown, the optimal monomer detachment rate $\delta_{opt}$. So, we need another equation, namely by minimizing the first-passage time. The mean first-passage time for the above random walk is given by

$$\langle T \rangle = \frac{L}{v} - \frac{D}{v^2}(1 - e^{-vL/D}). \tag{8}$$

What is left to do is to determine $m$, $K$, $\langle l \rangle$ and $\delta_{opt}$ self-consistently from 5, 6 and 7 and from minimizing the mean first-passage time 8.

As a first step, we use condition 7 to write $\delta = \nu m + \mu \frac{m^2}{K}$. Correspondingly, we find

$$D = 2\nu m + \mu \frac{m^2}{K}$$
$$v = -2\mu \frac{m^2}{K}$$

for the drift and diffusion constant in terms of $m$ and $K$. Using condition 6 together with the particle conservation condition 5 and with the mean-first passage time 8, we end up with the two defining equations for $m$ and $K$:

$$N = m + \frac{\nu}{\mu}\frac{K^2}{m} + \frac{K}{2} + \frac{S^2 K}{2(S + (\frac{\nu}{\mu}\frac{K}{m} + \frac{1}{2})(1 - e^{\frac{Sm}{\frac{\nu}{\mu}K + \frac{m}{2}}}))}$$
$$\langle T \rangle = -\frac{K}{2\mu m^2}(S + (\frac{\nu}{\mu}\frac{K}{m} + \frac{1}{2})(1 - e^{\frac{Sm}{\frac{\nu}{\mu}K + \frac{m}{2}}})).$$

To make progress, we make a last approximation, namely that $m \ll K$. This assumption is justified a posteriori and leads to

$$N = \frac{\nu}{\mu}\frac{K^2}{m} + \frac{S^2 K}{2(L + \frac{\nu}{\mu}\frac{K}{m}(1 - e^{S\frac{\mu m}{\nu K}}))}$$
$$\langle T \rangle = -\frac{K}{2\mu m^2}(S + \frac{\nu}{\mu}\frac{K}{m}(1 - e^{S\frac{\mu m}{\nu K}}))$$

or, in slightly rewritten form,

$$\frac{(S\frac{\mu m}{\nu K})^2}{2(1 - \frac{\mu m N}{\nu K^2})} = e^{S\frac{\mu m}{\nu K}} - 1 - S\frac{\mu m}{\nu K} \tag{9}$$
$$\langle T \rangle = \frac{S^2 K^2}{4\mu m(\frac{\nu}{\mu}K^2 - Nm)}. \tag{10}$$

Intriguingly, the first condition 9 is recast in terms of two dimensionless variables $a = \frac{S\mu m}{\nu K}$ and $b = \frac{N\mu m}{\nu K^2}$ as

$$e^a - 1 - a = \frac{1}{2(1-b)}a^2 \tag{11}$$

whose possible solutions are independent of all other parameters of the system and, in particular, independent of $S$. Furthermore, the average first-passage time then becomes

$$\langle T \rangle = \frac{\mu}{\nu^2}\frac{S^4}{4N}\frac{b}{a^2(1-b)}. \tag{12}$$

In order to minimize $\langle T \rangle$, thus, the term $b/(a^2(1-b))$ has to be minimized under the constraint 11. This minimization procedure is entirely independent of $S$ and we conclude that the average first-passage time scales as

$$\langle T \rangle \sim \frac{\mu S^4}{4\nu^2 N}. \tag{13}$$

Similarly, $m$ and $K$ behave as

$$m = \frac{\nu}{\mu} \frac{N}{S^2} \frac{a_{opt}^2}{b_{opt}} \sim \frac{\nu N}{\mu S^2}$$
$$K = \frac{N}{S} \frac{a_{opt}}{b_{opt}} \sim \frac{N}{S}.$$

From these scaling functions, we can finally determine the scaling of $\delta_{opt}$ from 7:

$$\delta_{opt} = \frac{\nu^2}{\mu}(m + \frac{m^2}{K}) = \frac{\nu^2}{\mu}(\frac{N}{S^2} \frac{a_{opt}^2}{b_{opt}} + \frac{N}{S^3} \frac{a_{opt}^3}{b_{opt}}) \sim \frac{\nu^2}{\mu} \frac{N}{S^2},$$

where we neglected the higher-order scaling $\sim \frac{N}{S^3}$. This yields the parameter exponent $\phi = -2$.

As a last step, we can actually determine $a_{opt}$ and $b_{opt}$ numerically from minimizing $b/(a^2(1-b))$ under the constraint 11. This procedure yields

$$a_{opt} \approx 2.687$$

$$b_{opt} \approx 0.672$$

and plugging in these values into the formulas for $\langle T \rangle$ and $\delta_{opt}$ we get:

$$\langle T \rangle \approx 0.07 \frac{\mu S^4}{\nu^2 N} \tag{14}$$
$$\delta_{opt} \approx \frac{\nu^2}{\mu}(10.74 \frac{N}{S^2} + 28.87 \frac{N}{S^3}) \approx 10.74 \frac{\nu^2}{\mu} \frac{N}{S^2}. \tag{15}$$

Combining the scaling behavior of $m$ and $\delta_{opt}$, we find that the drift coefficient $D$ vanishes to lowest order and the polymer size distribution behaves as a purely diffusive process. Intriguingly, this is true not only in the optimal case but follows more generally from the quasi-stationarity assumption: the system self-organizes into a diffusion process without drift where growth of structures and detachment of monomers balance. The optimal parameter choice thus corresponds to maximizing the diffusive flux through the system.

**Irreversible scenarios and reversible binding for 2D/3D structures**

The simulations show that in the irreversible scenarios, the respective control parameter is optimal (achieving a minimal $T_{90}$ assembly time) close to where the *final* yield is approximately 90% (see main text Fig. 3C,D). This is plausible because in all scenarios the control parameter defines the rate limiting time scale and hence the parameter is optimal close to where the desired yield is barely reached. Therefore, the scaling of the optimal parameter can be determined by identifying a scaling relation that fixes a constant final yield. In order for the final yield to be independent of the size $S$ of the target structure, the ratio between the total nucleation and total attachment rate must scale inversely with $S$. To put it more simply: if the size of the target structure is doubled, in order to achieve a constant yield, there need to be twice as many growth events relative to the same number of initiation events.

$$\frac{\text{total number of nucleation events per time}}{\text{total number of attached monomers per time}} := \frac{\mu_{\text{tot}}}{\nu_{\text{tot}}} \overset{!}{\sim} \frac{1}{S} \tag{16}$$

This formula provides the starting point of our argument. In the following paragraphs we identify the total nucleation rate $\mu_{\text{tot}}$ and total attachment rate $\nu_{\text{tot}}$ for the three irreversible scenarios as well as for the reversible binding scenario in higher dimensions.

**Activation and dimerization scenario**

In the activation and dimerization scenario, we focus on one-dimensional structures only. The higher dimensional cases are related to the one-dimensional case via the scaling argument provided in the main text.
The total nucleation rate depends quadratically on the momentary concentration of active monomers $m$ per species and linearly on $S$ (number of possible dimerization partners).

$$\mu_{\text{tot}} = \mu m^2 S \tag{17}$$

The total attachment rate is given by the product of the total concentration of complexes $K$ in the system and the concentration of monomers per species.

$$\nu_{\text{tot}} = \nu K m \tag{18}$$

Note that the total concentration of complexes $K$ will scale with $C = \frac{N}{V}$ but can be assumed to be independent of $S$ as we demand a constant yield. Therefore,

$$\frac{\mu_{\text{tot}}}{\nu_{\text{tot}}} \sim \frac{\mu S m}{\nu C} \overset{!}{\sim} \frac{1}{S}. \tag{19}$$

In the dimerization scenario, all particles are active from the outset, hence $m \sim C$ and therefore, $\mu^{\text{opt}} \sim \frac{\nu}{S^2}$. Because dimerization is the time-limiting process in the dimerization scenario, this implies for the minimal assembly time

$$T_{90}^{\text{min}} \sim \frac{C}{\mu_{\text{tot}}^{\text{opt}}} \sim \frac{1}{SC\mu^{\text{opt}}} \sim \frac{S}{C\nu}. \tag{20}$$

So, the argument reproduces the control parameter exponent $\phi = -2$ and the time complexity exponent $\theta = 1$ for the dimerization scenario for one-dimensional structures. The dependence on $\nu$ is crucial for the rescaling argument described in the main text in order to relate to the higher dimensional cases.
In the activation scenario, the monomers are not active right from the outset. Instead, there is a constant influx of monomers that balances a steady consumption of monomers due to binding. Hence, the stationary concentration $m$ of active monomers is determined from the condition that the total influx of monomers equals their consumption due to binding:

$$\text{total influx rate of monomers} = \text{total consumption of monomers due to binding.} \tag{21}$$

With the total influx rate of monomers given by $\alpha CS$ this translates into

$$\alpha CS = \nu_{\text{tot}} \sim \nu K m, \tag{22}$$

where we neglected the consumption of monomers due to dimerization because for large $S$ dimerization is negligible compared to attachment (compare Eq. (16)). With $K \sim C$ this gives $m \sim S\frac{\alpha}{\nu}$ and hence with Eq. (19),

$$\alpha^{\text{opt}} \sim \frac{\nu^2 C}{\mu} \frac{1}{S^3} \ . \tag{23}$$

Furthermore, because the influx rate limits the assembly time,

$$T_{90}^{\text{min}} \sim \frac{1}{\alpha^{\text{opt}}} \sim \frac{\mu}{C\nu^2} S^3 \ , \tag{24}$$

confirming the control parameter exponent $\phi = -3$ and time complexity exponent $\theta = 3$ for the one-dimensional activation scenario, as well as a quadratic dependence on $\nu$ that is relevant for the rescaling procedure to relate to the higher dimensional cases.

**JIS scenario**

In the JIS scenario the different species are provided sequentially in consecutive batches. In order to estimate the total nucleation and attachment rate in Eq. (16), we calculate the total number of nucleation and binding events per species in a single assembly step. The number of nucleation events will crucially be determined by the number of active monomers that are still unbound when the next batch is supplied: Since the subsequent batch is supplied when most monomers of the previous batch have already bound, the remaining monomers encounter many partners to form dimers while there are only few remaining binding sites in the clusters. Therefore, the remaining monomers will dimerize to the largest extent. In order

to estimate the total dimerization rate per assembly step, we therefore need to estimate the concentration of remaining monomers in relation to $\Delta T$. We do this in the following by considering the dynamics of the concentration of monomers of an arbitrary species in the sequence.
Let $m$ denote the concentration of monomers of a species $i$ and $k$ the concentration of structures (binding sites) to which species $i$ can attach. We assume that at time $t = 0$, species $i$ is supplied in initial concentration $m(0) = M \sim C$. Each binding event involving species $i$ reduces both the concentration of binding sites $k$ and the concentration of monomers $m$ by one unit. Therefore, $u := m - k = \text{const}$ is a constant which denotes the excess concentration (i.e. the amount by which the total number of monomers $M$ exceeds the number of binding sites $k(0) := K$). Indeed, $u$ corresponds to the increase in concentration from one batch to the next in the context of non-stoichiometric concentrations (see Methods). For the dynamics of $m$ it then follows that

$$\frac{d}{dt}m = -\nu mk = -\nu m^2 + \nu um. \tag{25}$$

By solving the differential equation we find the monomer concentration $m$ at time $t = \Delta T$:

$$m(\Delta T) = \frac{1}{\frac{1}{u} + \left(\frac{1}{M} - \frac{1}{u}\right) e^{-u\nu\Delta T}} \approx \frac{1}{\frac{1}{u} + \left(\frac{1}{M} - \frac{1}{u}\right)(1 - u\nu\Delta T)} \approx \frac{1}{\nu\Delta T}, \tag{26}$$

where in the second step we assumed $\Delta T \sim 1/(C\nu) \ll 1/(u\nu)$ and in the last step we used $1/M \ll 1/u$ (or $u \ll M$). The total number of dimerization events during one assembly step can now be estimated as the concentration of monomers of species $i$ that are still unbound at time $\Delta T$ when the next binding partner, species $i + 1$, is supplied (in concentration $\approx M$). More specifically, the total number of dimerization events per assembly step is $\sim m(\Delta T)M \sim \mu_{\text{tot}}$ while the total number of attachment events per assembly step is $\sim KM \sim \nu_{\text{tot}}$ where $K := k(0) \sim C$. Therefore, with Eq. (16),

$$\frac{\mu_{\text{tot}}}{\nu_{\text{tot}}} \sim \frac{1}{\nu C\Delta T} \overset{!}{\sim} \frac{1}{S}. \tag{27}$$

and thus,

$$\Delta T^{\text{opt}} \sim \frac{S}{C\nu}, \tag{28}$$

yielding the control parameter exponent $\phi = 1$. In order to obtain the total assembly time, $\Delta T$ must be multiplied by the total number of batches, which is $b_{\text{max}} \sim S^{1/d}$ in the case of the 'onion-skin' supply protocol (see main text). Therefore,

$$T_{90}^{\text{min}} \sim \Delta T^{\text{opt}}\ S^{1/d} \sim \frac{S^{1+\frac{1}{d}}}{C\nu}, \tag{29}$$

yielding the time complexity exponent $\theta = 1 + \frac{1}{d}$, where $d$ is the dimensionality.

#### Reversible binding for 2D and 3D structures

For the reversible binding scenario in two and three dimensions we can use the same approach as for the irreversible scenarios, starting from Eq. (16). The key insight is that during the assembly process stable intermediate assembly products form that decay only with rate $\delta_2 \ll \delta_1$ or $\delta_3 \lll \delta_1$ and hence are considered as long-lived on the relevant timescale. In contrast, intermediate states that decay with rate $\delta_1$ are highly unstable and decay quickly as $\delta_1$ is typically large compared to the reactive timescale $C\nu$ in the reversible binding scenario. Figure S4 shows how the resulting total nucleation rate $\mu_{\text{tot}}$ ($\mu_{\text{tot}}$ here denotes the total nucleation rather than dimerization rate) and total attachment rate $\nu_{\text{tot}}$ can be estimated. Here nucleation is an effective four-particle reaction that proceeds via two unstable intermediate states. If the detachment rate $\delta_1$ is large, the effective per capita rate for the four-particle reaction can be approximated as $\mu\nu^2/\delta_1^2$ and the total nucleation rate is given by $\mu_{\text{tot}} \sim \frac{\mu\nu^2}{\delta_1^2} m^4 S$ (the factor $S$ accounts for the fact that there are S possible combinations of species that can form a nucleus). Attachment typically proceeds in two steps. The first step, analogous to the nucleation process, can be approximated as an effective two-particle reaction passing through an unstable intermediate state (see Figure S4). The effective total rate for the first step is therefore $\sim \frac{\nu^2}{\delta_1^2} K m^2 S^{\frac{d-1}{d}}$, where $K$ is the total concentration of complexes and the factor $S^{\frac{d-1}{d}}$ estimates the number of possible binding sites for the first monomer (surface area of an average cluster). Once a new stable state has formed, a cascade of subsequent stable states can be traversed by attachment of additional monomers. Because in this second step the complex only passes through stable states, the second step can be assumed to be fast compared to the first step. We estimate the average number of monomers attaching in the second step to scale again proportionally to the cluster surface $\sim S^{\frac{d-1}{d}}$. This yields an additional stoichiometric factor to be accounted for in the total attachment rate, resulting in $\nu_{\text{tot}} \sim \frac{\nu^2}{\delta_1} K m^2 S^{(2-\frac{2}{d})}$. With Eq. (16) it follows that

$$\frac{\mu_{\text{tot}}}{\nu_{\text{tot}}} \sim \frac{\mu C}{\delta_1} S^{(\frac{2}{d}-1)} \overset{!}{\sim} \frac{1}{S} , \tag{30}$$

and, therefore,

$$\delta_1^{\text{opt}} \sim \mu C S^{\frac{2}{d}}, \tag{31}$$

with a control parameter exponent $\phi = \frac{2}{d}$. Since nucleation is the slowest step, we expect the minimal assembly time to scale approximately as the timescale of nucleation:

$$T_{90}^{\text{min}} \sim \frac{C}{\mu_{\text{tot}}} \sim \frac{C}{\mu\left(\frac{\nu}{\delta_1^{\text{opt}}}\right)^2 m^4 S} \sim \frac{\mu}{\nu(C\nu)} S^{\frac{4}{d}-1} , \tag{32}$$

yielding a time complexity exponent $\theta = \frac{4}{d} - 1$. Although the theoretical estimates for the exponents in the reversible binding scenario in higher dimensions do not coincide perfectly with the simulated values (compare main text Fig. 2B,C and Fig. 3B), their tendency and the dependence on the dimensionality of the structure are correctly predicted and explained. We suspect that the main reason for the deviations is a slight actual dependence of the average monomer concentration $m$ on $S$, which has been neglected in this scaling argument.

In conclusion, note that for all four scenarios, the scaling exponents for one-dimensional structures could be derived exactly from our scaling analysis. In contrast, for higher dimensional structures, the theoretical estimates generally do not fit the simulated values exactly. This may have various reasons like, for example, deviations from the presumed effective growth rate $\nu_S \sim \nu S^{\frac{d-1}{d}}$ that we used to rescale the exponent for the dimerization and activation scenario (see main text).
Furthermore, note that in the scaling argument applied to the reversible binding scenario in higher dimensions we used some specificities of the structure, most importantly, the number of unstable intermediate states in the processes of nucleation and attachment. This suggests that the exponents and the time efficiency of the reversible binding scenario are not fully generic but depend on the shape of the structure and the constituents. In contrast, the scaling arguments for the other scenarios are fully generic, so we do not expect a significant dependence of the time efficiency on specificities of the structure in the irreversible scenarios.

### 3. ROBUSTNESS TO MODEL MODIFICATIONS

To verify that our time complexity analysis is robust to model modifications, we investigated three variants of the original model and the assembly kinetics. Figure S5 shows the minimal assembly time for these variants in all four scenarios. The results of the analysis are discussed in the following.

#### Structures with periodic boundaries

First we simulated the minimal assembly time for structures with periodic boundaries. While in the main text we only considered structures with an open boundary, some typical examples of self-assembling systems comprise the formation of closed structures with periodic boundaries, for instance, one-dimensional rings [2] or two-dimensional shells and capsids such as, for example, virus capsids [3, 4]. To assess the relevance of the boundary we simulated the minimal assembly time for two-dimensional periodic structures or tori. In all scenarios we measured almost the same time complexity exponent as in the original model. Only in the reversible binding scenario the exponent appears to be slightly larger. In the activation, dimerization and reversible binding scenario, the time efficiency increases as a consequence of the modified boundary condition since a closed boundary effectively enhances the possibility of a cluster to grow, thereby increasing the effective binding rate. In the JIS scenario, the time efficiency slightly decreases because the species at the boundary induce an increased excess dimerization rate compared to the case with non-periodic boundary. Also note that, in the JIS scenario, we simulated periodic structures only with an even edge length $L$, since for odd $L$ it would have been necessary to modify the supply order of our protocol in order to make sure that species supplied in the same batch do not bind each other. Moreover, we increased the excess concentrations $E_n$ (see Methods) of the species at the boundary by a factor of 2 or 4, respectively, to achieve optimal efficiency for the modified boundaries.

#### Heterogeneous binding rates

Next, we investigated the impact of heterogeneous binding rates on the assembly time. Considering a heterogeneous system, the assumption of identical binding rates for all species is an idealization. More realistically, the rates will vary to a certain extent. We therefore simulated the system with heterogeneous rates for the different species, drawn independently from a (truncated) normal distribution with a coefficient of variation of 50%. We truncated the normal distribution for values that are below 20% of the mean in order to guarantee that individual rates do not become negative or very small. For each run the binding rates were chosen independently and the assembly times were averaged over 10-100 independent runs. We did not perform the simulations for the activation scenario since the simulation of the activation scenario is based on the homogeneous approximation and the results would thus not be reliable for heterogeneous rates of the species. In the other scenarios, the measured time complexity exponents are almost identical to those of the original model with homogeneous rates. Only in the dimerization scenario the time complexity exponent seems somewhat smaller, probably because heterogeneity in the rates influences the typical shapes in which clusters grow. In all cases, the time efficiency was reduced as a consequence of heterogeneous rates because small rates influence the overall effective timescale more significantly than the large rates.

**Reduced resource efficiency**

Finally, we altered the definition of the assembly time and explored its effect on the time complexity. In the main text we chose 90% yield as termination criterion for the assembly process. Here, we asked whether the exponents are invariant if a lower resource efficiency of only 50% yield is demanded. In all scenarios, the minimal time $T_{50}^{\min}$ required to achieve 50% yield is significantly smaller than $T_{90}^{\min}$. With the exception of the activation scenario, however, the corresponding time complexity exponents are indistinguishable from those determined for $T_{90}^{\min}$. For the activation scenario, the exponent appeared to be a bit larger, very close to the theoretical value $\Theta_{\mathrm{th}} = 2$. We relate this slight discrepancy between the two exponents to the fact that the yield transition curves in the activation scenario (compare main text Fig. 3D) become steeper if S is increased. This indicates that the real asymptotic exponent for the activation scenario lies between the exponents measured for $T_{50}^{\min}$ and $T_{90}^{\min}$. Note that, among the four scenarios, the time efficiency of the reversible binding scenario increases the most if lower resource efficiency is demanded.

**Annealing (reversible binding scenario)**

The reversible binding scenario is controlled by the ratio between the detachment rate and the growth rate, given by the product of the binding rate $\nu$ and the concentration of monomers (see main text, paragraph reversible binding scenario). However, when more and more particles get attached during the assembly process, the concentration of monomers - if not replenished - gradually decreases. Consequently, the controlling parameter increases during the assembly process. In order to counteract this effect, a frequently used experimental approach consists in 'annealing' the system by decreasing the temperature [5]. Typically, one starts at high temperature and gradually cools the system down to room temperature. Since the detachment rate decreases with decreasing temperature, $\delta \sim e^{-E_B/(k_B T)}$, if applied optimally, annealing allows to keep the ratio between detachment rate and growth rate constant during the assembly process. Here we ask how the time efficiency in the reversible binding scenario behaves under an optimal annealing protocol. To this end, we assume that the temperature adapts instantaneously to the momentary concentration of monomers such that the ratio between detachment rate and monomer concentration remains constant throughout the simulation. By varying this fixed ratio we determine the minimal assembly time $T_{90}^{\min}$ as in the main text. Indeed we find that the assembly efficiency can be significantly increased with an optimal annealing protocol, however, the time complexity exponent remains invariant (see Figure S5A, star marker).

**Alternate input functions (activation scenario)**

For the activation scenario in the main text we assumed a constant influx of active monomers until all inactive monomers are depleted. Hence, the input as a function of time has a rectangular shape. A natural question that arises is whether the efficacy of the activation scenario can be altered by changing the temporal form of the input. To answer this question we simulated various different input functions which correspond to different biophysical processes providing the active monomers. Here we discuss one particular example for such a differing form of the temporal input which plays an important role in biology [6]. Specifically, we assume that activation of monomers is no longer irreversible but, instead, monomers can switch back and forth between an assembly-active and inactive configuration (reversible activation cycle). Furthermore, we assume that this switching dynamics is fast compared to the assembly time scale and hence can be considered to be at equilibrium. The control parameter is the equilibrium constant $K$, which describes the ratio between the concentrations of active and inactive monomers. By measuring the minimal assembly time in the usual way, we find that the activation scenario becomes slightly more efficient through the reversible activation cycle but that the time complexity exponent remains invariant (see Fig. S5C, star marker).
Theoretically, the input can be described by any arbitrary function that integrates to the total particle number $N$. Note that input via fast reversible activation has a special significance because through equilibration it allows the net influx rate to dynamically adopt to the current state of the assembling system (fast binding of active monomers $\rightarrow$ fast net influx, and vice versa). We also tested some other input functions and observed that it generally seems to be favourable for the time efficiency if the input is higher at the beginning of the assembly process and lower towards the end. The measured time complexity exponents however remained invariant for all tested input functions. This leads us to hypothesise that the time complexity exponent cannot be altered by the form the monomer input as long as all species are treated indifferently.

In conclusion, we tested how robust our results are with respect to modifications of the model, affecting the boundary of the structures, heterogeneities in the rates or the demanded resource efficiency. Furthermore, we investigated differing experimental protocols like annealing or variable input functions for the activation scenario. We found that while the assembly time does indeed depend on details of the model and the assembly protocol, the time complexity exponents - apart from minor deviations - remain invariant to such variations. Furthermore, the general trend in response to a particular model variation is typically the same in the different scenarios (an exception is the modification of the boundary condition in the JIS scenario). This confirms that the general conclusions in the main text on the time efficiency of the different scenarios and their relative ranking remain largely valid if details of the system are changed. On a broader perspective, this shows that the time complexity analysis yields a reliable, robust and informative characterization of self-assembly processes and the distinction of the four scenarios, characterized by different time complexity exponents, is meaningful and useful.

### 4. EXPERIMENTAL JIS SUPPLY PROTOCOL FOR THE ASSEMBLY OF AN ARTIFICIAL T=1 CAPSID

In this last chapter we aim to demonstrate the applicability of the just-in-sequence supply strategy for actual experimental problems of interest by proposing a specific supply protocol for the assembly of an artificial $T = 1$ capsid.

Artificial shells and capsids have important potential biotechnological applications ranging from the compartmentalization of chemical reactions to the usage as vesicles that enable pinpoint delivery of drugs or other material to specific loci within an organism [7, 8]. The simplest icosahedral capsid is the $T = 1$ capsid (classification by Caspar and Klug, [9] which is assembled from 60 proteins. In the following, we discuss two possibilities to assemble artificial T=1 capsids irreversibly with high yield solely by regulating the supply of constituents. These strategies thereby avoid the necessity of fine-tuning the binding strengths or other molecular properties. The first possibility assumes a partly homogeneous design of the capsid (see Fig. S6A), while the second possibility relies on a fully heterogeneous design (Fig. S6C) of the structure.

In principle, the $T = 1$ capsid can be build fully homogeneously out of 60 identical units. However, in order to use the just-in-sequence supply strategy as described in the main text, some degree of heterogeneity is necessary: constituents that are provided in the same batch should not be able to bind each other but only to the existing structures. We therefore propose the partly heterogeneous design depicted in Fig. S6A, which exploits the symmetry of the target structure. Components that are indicated by the same letter are identical and bind specifically only with those species that are adjacent to them.
Designing structures as homogeneously as possible has three practical advantages. First, a lower number of different components needs to be produced and counted, which reduces the experimental effort. Second, self-assembly is faster if a single type of constituent can bind to several distinct sites in the structure and finally, as we discuss below, the absolute tolerance to external noise in particle numbers increases if structures are more homogeneous.
Note, however, that for the assembly of spherical objects like the T=1 capsid, a difficulty arises concerning the upper and the lower ”cap“, denoted here by A and L, respectively: If the caps are composed of several copies of a single species, these copies would be able to form homo-multimers when they are supplied, thereby undermining the JIS strategy. This challenge can be circumvented either by designing the caps heterogeneously or by making the respective bonds between the cap-species weak and reversible, thereby preventing spurious nucleation. Another possibility is to produce the caps A and L separately and supply them as single, complete units. In the following, for the assembly of the partly homogeneous capsid, we further discuss the second possibility, considering the caps A and L as single units.

Figures S6B and D show possible supply protocols for the assembly of the partly homogeneous and the heterogeneous T=1 capsid, respectively. Both of these protocols were found by maximizing the yield in the simulation. The second column in the tables indicates the species that are supplied in the respective batch, while the third column shows the numbers $Z_b$ that describe the excess concentrations supplied for the species in the respective batch, see Methods. The total number $N_b$ of particles for each species supplied in the $b^{\text{th}}$ batch (fourth column) is given by (compare Methods)

$$N_b = \deg \cdot \left( (1-p)N + pSN \frac{Z_b}{Z_{\text{tot}}} \right) , \tag{33}$$

where deg is the *degeneracy* of the species, denoting the number of distinct binding positions per structure for this species in the respective assembly step. For the partly homogeneous capsid, the degeneracy is deg = 5 for all species except for the caps which are provided as complete units with degeneracy deg = 1. It is likely that the efficiency of the supply strategy can be further improved by allowing the pairs of species C and D, F and G, as well as I and J, which are supplied in the same batches, to be assigned different particle numbers. For simplicity, however, in this example we assign particle numbers only in correspondence to the batch number.

Figure S7A shows the yield plotted against the interval $\Delta T$ between successive batches both for the partly homogeneous and the heterogeneous $T = 1$ capsid. Black circles indicate the position of the optimal interval $\Delta T_{\text{opt}}$ that minimizes the time required to achieve 90% yield. The partly homogeneous capsids can be assembled in shorter time (provided that the same number of structures is assembled) because the binding speed is larger roughly by a factor of 5 compared to the fully heterogeneous $T = 1$ capsid.

In applications, particle numbers can only be determined with limited accuracy. Hence, it is an essential question how robust this approach is to extrinsic noise in the particle numbers. In order to test the robustness to extrinsic noise we choose particle numbers randomly from a Gaussian distribution and quantify the noise level in terms of the coefficient of variation (CV), defined as the standard deviation of the particle numbers relative to their respective mean. For simplicity, we assume that the CV is the same for all species. Figure S7B shows the yield plotted against the time interval $\Delta T$ for the partly homogeneous $T = 1$ capsid depending on the coefficient of variation. The inset shows the maximum yield (achieved for sufficiently large $\Delta T$) plotted against the CV, both for the partly homogeneous and the heterogeneous design. As a rough estimate, for the two supply protocols discussed here, particle numbers would need to be chosen with an accuracy of about 1% in order to achieve high yield. For a fixed relative strength of noise compared to the mean (CV), the partly homogeneous capsid is slightly more robust than the heterogeneous structure. This implies that the absolute tolerable variability in the number of particles per species is larger by at least a factor of 5 for the partly homogenous capsid compared to the heterogeneous capsid.

In conclusion, we found that both the partly homogeneously as well as the heterogeneously designed $T = 1$ capsid could be assembled efficiently with an irreversible just-in-sequence supply strategy provided that particle numbers can be determined accurately enough. The supply protocols discussed here still leave space for improvement, for example, by assigning particle numbers individually for each species rather than only in correspondence to the batch number. Furthermore, the excess concentrations were chosen in order to guarantee maximal yield for $\Delta T \to \infty$ but have not been optimized for maximal robustness to external noise. Those improvements might allow to even further improve the efficiency and robustness of the approach. Hence, provided that experimental methods for the accurate counting of molecules can be established, the JIS scenario offers a versatile strategy for the realization of biotechnologically relevant macromolecular structures. Our work therefore highlights how new experimental strategies to control concentrations could advance nanotechnology and its applications.

### B. Supplementary Figures

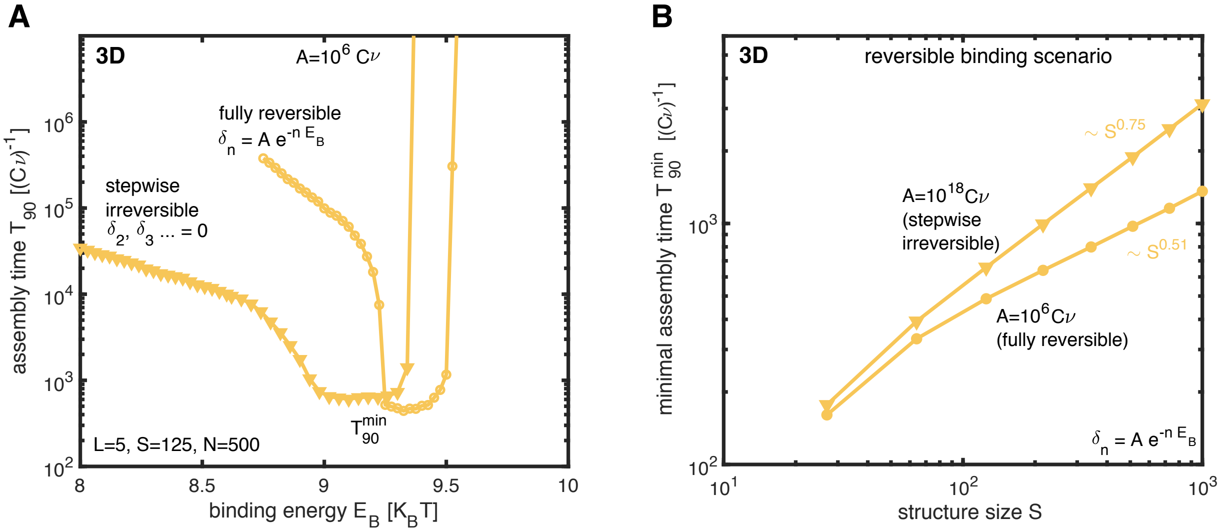


**Figure S1: Reversible binding scenario: influence of the preexponential factor on the assembly time. A**, assembly time $T_{90}$ versus the binding energy $E_B$ for small preexponential factor $A = 10^6 C\nu$ (marker: circle) for three-dimensional structures of size $S = 125$. For comparison, we also plotted the stepwise irreversible case (marker: triangle) setting all detachment rates except for $\delta_1$ to 0. The stepwise irreversible case is equivalent to choosing $A$ large (in the main text: $A = 10^{18} C\nu$) as in both cases only $\delta_1$ is effectively larger than 0 and all other detachment rates are negligible at close-to-optimal binding energies. Hence, a small preexponential factor A slightly decreases the minimal assembly time (compared to large $A$) at the cost of a reduced variability in the binding energy (fine tuning of $E_B$ (or of the concentration $C$) becomes more critical with small $A$). **B**, minimal assembly time $T_{90}^{\text{min}}$ versus the structure size $S$ for large (stepwise irreversible) and small (fully reversible) preexponential factor $A$. The minimal assembly time that can be achieved as well as the time complexity exponent are slightly smaller for a small preexponential factor.

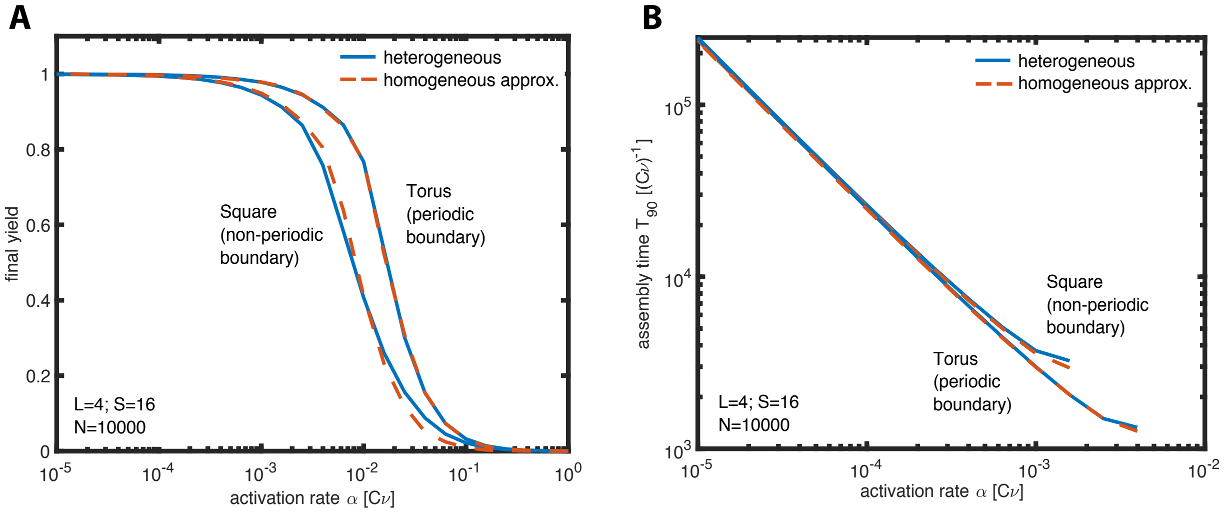


**Figure S2: Accuracy of the homogeneous approximation in the activation scenario.** Final yield (**A**) and assembly time $T_{90}$ (**B**) versus the activation rate. Both quantities were simulated for two-dimensional structures with and without periodic boundaries as well as with distinguishable (heterogeneous, blue drawn line) and indistinguishable particle species (homogeneous approximation, red dashed line). For structures with periodic boundaries and large particle number $N$, the homogeneous and heterogeneous simulation coincide exactly as predicted by the theory (see SI). For structures with non-periodic boundaries, the homogeneous system yields an accurate approximation of the heterogeneous system. We exploited this equivalence to reduce the computational cost by simulating the activation scenario as a homogeneous system with lower particle number. Generally, the heterogeneous system is subject to stochastic effects arising from fluctuations between the concentrations of the different species, unless the particle number $N$ is large (see reference [1]). The homogeneous system, in contrast, can be simulated with a much smaller total number of particles.

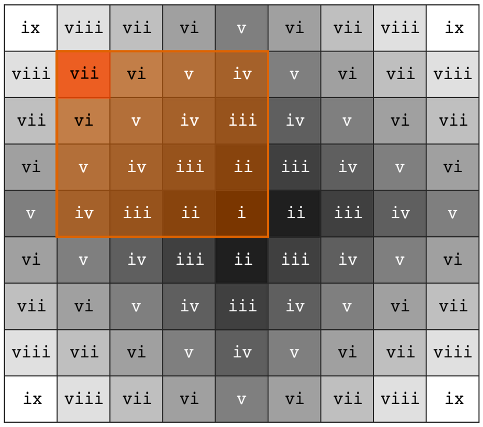


**Figure S3: Assigning particle numbers in the Jis scenario.** The just-in-sequence scenario requires specified ratios between particle numbers in order to avoid excessive competition for resources (see main text and methods). Shown is the onion supply protocol (analogous to Fig. 4C) for a two-dimensional structure of size L=9 (S=81). Roman numbers indicate the batch number (assembly step) in which species are supplied. The shaded square marks all species that can initiate a complex potentially able to bind the species highlighted in red in the seventh assembly step. In order to minimize competition for resources, the species in the seventh batch must hence be supplied in excess concentration $Z_7$ proportional to the area of the square to allow all clusters present at the seventh assembly step to grow. Generalizing, we hence find the excess concentration $Z_n \sim \left(\frac{(n+1)}{2}\right)^2$ for a species supplied in the $n^{\text{th}}$ batch (compare Methods).

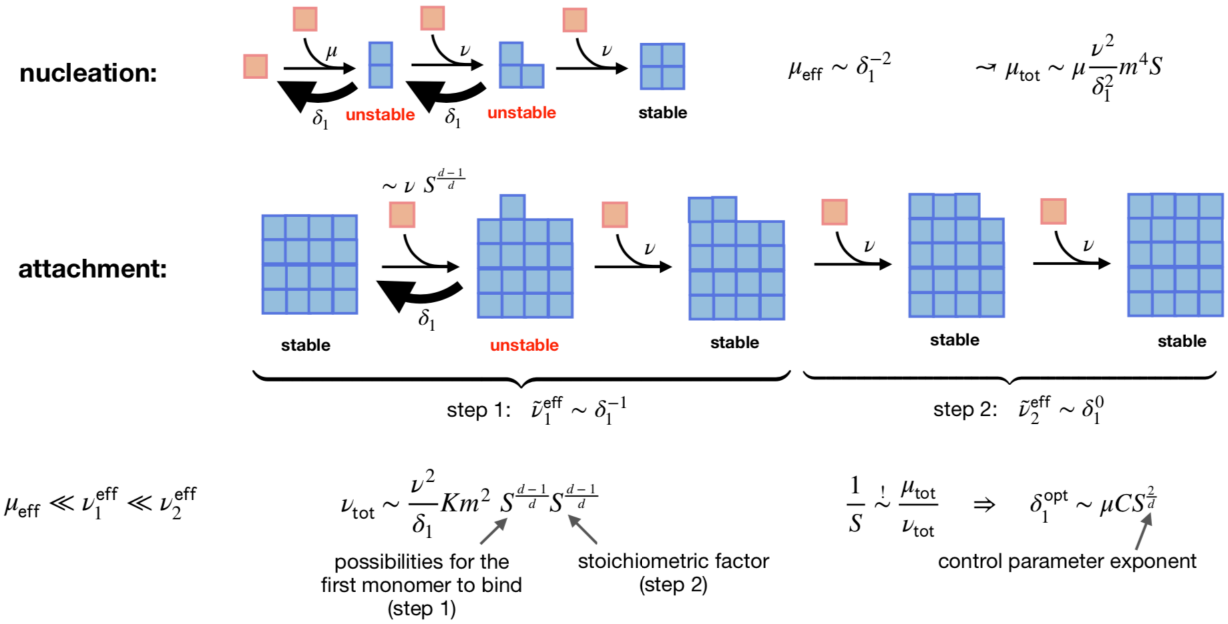


**Figure S4: Scaling analysis of the reversible binding scenario.** In the reversible binding scenario, a stable nucleus forms by passing through two unstable intermediate states that decay with rate $\delta_1$. Hence, the effective rate for the nucleation process is $\mu_{\rm eff} \sim \delta_1^2$. Attachment typically proceeds in two steps. In the first step, a monomer first binds reversibly and must subsequently be stabilized by a second monomer. Because one unstable state is passed, the first step effectively happens at rate $\nu_1^{\rm eff} \sim \delta_1^1$. Subsequently to the first step, additional monomers can attach 'filling the row', while the configuration is continuously stable. Therefore, the second step can be assumed to be fast compared to the first step which, in turn, is fast compared to nucleation: $\mu_{\rm eff} \ll \nu_1^{\rm eff} \ll \nu_2^{\rm eff}$. By setting the total nucleation rate into relation with the total effective attachment rate as detailed in Supplement section 3, a rough estimate for the control parameter exponent and for the time complexity exponent can be derived.

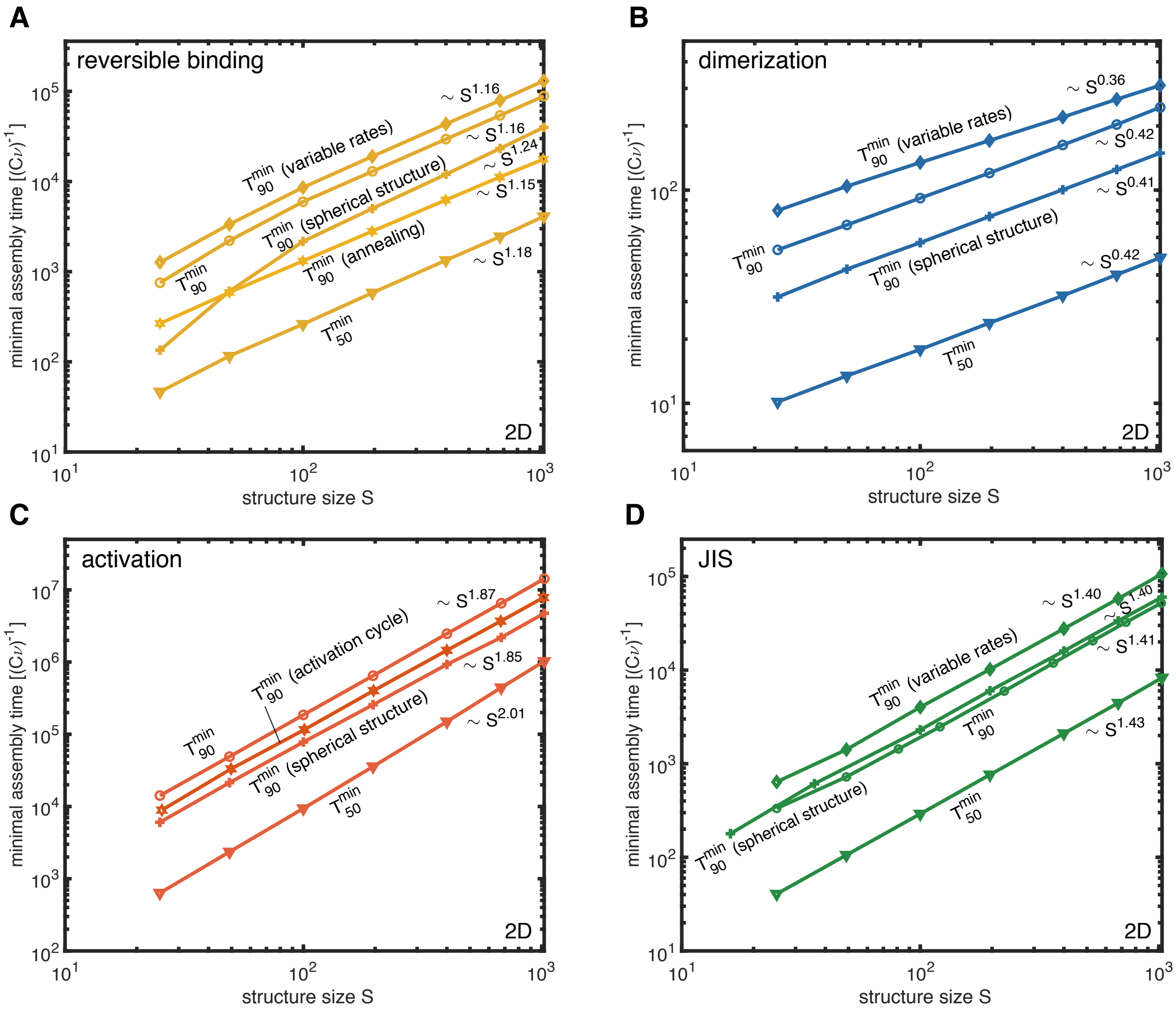


**Figure S5: Scaling of the minimal assembly time for variants of the model and assembly kinetics.** The minimal time required to achieve 90% ($T_{90}^{\mathrm{min}}$) or 50% yield ($T_{50}^{\mathrm{min}}$) in the different scenarios (**A,** reversible binding; **B,** dimerization; **C,** activation and **D,** just-in-sequence scenario) is shown in dependence of the target structure size $S$ for two-dimensional structures and different variants of the original model. In each subpanel (scenario), the curve labeled $T_{90}^{\mathrm{min}}$ corresponds to the assembly time in the original model. Furthermore, each subpanel shows $T_{90}^{\mathrm{min}}$ for 2D structures with periodic boundary (tori) as well as for variable or heterogeneous rates of the constituent species (not available for the activation scenario), see SI. The curve labelled $T_{50}^{\mathrm{min}}$ shows the minimal assembly time for a lower resource efficiency of only 50% yield. While the assembly time varies for the different model variants, the measured time complexity exponents are, aside from small deviations, (see SI) largely invariant. This indicates that the time complexity analysis of the self-assembly scenarios is robust and independent of many details of the model.

**A**

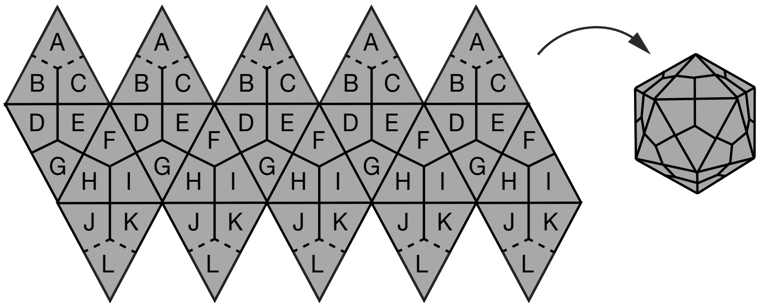


**B**

| batch number | supplied species | excess conc. $Z_b$ | particle number $N_b$ |
|---|---|---|---|
| 1 | A | 0 | 0.9N |
| 2 | B | 1 | 4.59N |
| 3 | C,D | 2 | 4.69N |
| 4 | E | 4 | 4.88N |
| 5 | F,G | 5 | 4.97N |
| 6 | H | 7 | 5.16N |
| 7 | I,J | 10 | 5.45N |
| 8 | K | 18 | 6.20N |
| 9 | L | 40 | 1.66N |

**C**

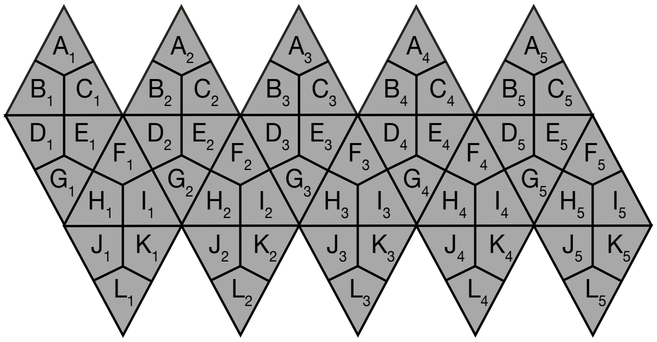


**D**

| batch number | supplied species | excess conc. $Z_b$ | particle number $N_b$ |
|---|---|---|---|
| 1 | $A_2$ | 0 | 0.930N |
| 2 | $A_1$, $A_3$ | 1 | 0.932N |
| 3 | $A_4$ | 3 | 0.935N |
| 4 | $A_5$ | 4 | 0.937N |
| 5 | B | 5 | 0.939N |
| 6 | C,D | 10 | 0.948N |
| 7 | E | 20 | 0.966N |
| 8 | F, G | 25 | 0.975N |
| 9 | H | 35 | 0.993N |
| 10 | I,J | 50 | 1.02N |
| 11 | K | 90 | 1.09N |
| 12 | $L_1$ | 95 | 1.10N |
| 13 | $L_2$, $L_5$ | 130 | 1.16N |
| 14 | $L_4$ | 170 | 1.24N |
| 15 | $L_3$ | 200 | 1.29N |

**Figure S6: Capsid structure and supply protocols.** **A**, Partly homogeneous design of the T=1 capsid consisting of 60 subunits and 12 different species. Species of subunits are indicated by capital letters. It is assumed that each species binds specifically only with those species adjacent to it. Furthermore, we assume that the caps, each consisting of 5 subunits of A and L, respectively, are assembled separately and are supplied as complete single units. **B**, Just-in-sequence supply protocol that was simulated in order to assemble the capsid with the structure defined in (A). Columns indicate the species that are supplied in a respective batch, their excess concentration and their resulting total particle numbers assuming a fraction of unevenly distributed resources of $p = 0.07$ (cf. Methods and Eq. (33). Here, $N$ is the number of complete structures to be built if the yield were 100%. **C**, Heterogeneous design of the T=1 capsid consisting of 60 subunits and 60 different species. Each species occupies a single specified position in the structure. **D**, Just-in-sequence supply protocol for the heterogeneous structure described in (B). Letters without indices in the protocol represent all 5 corresponding species (for example $B = \{B_1, B_2, B_3, B_4, B_5\}$), which are supplied simultaneously. Note that for the heterogeneously designed capsid the caps are assembled from monomers as well.

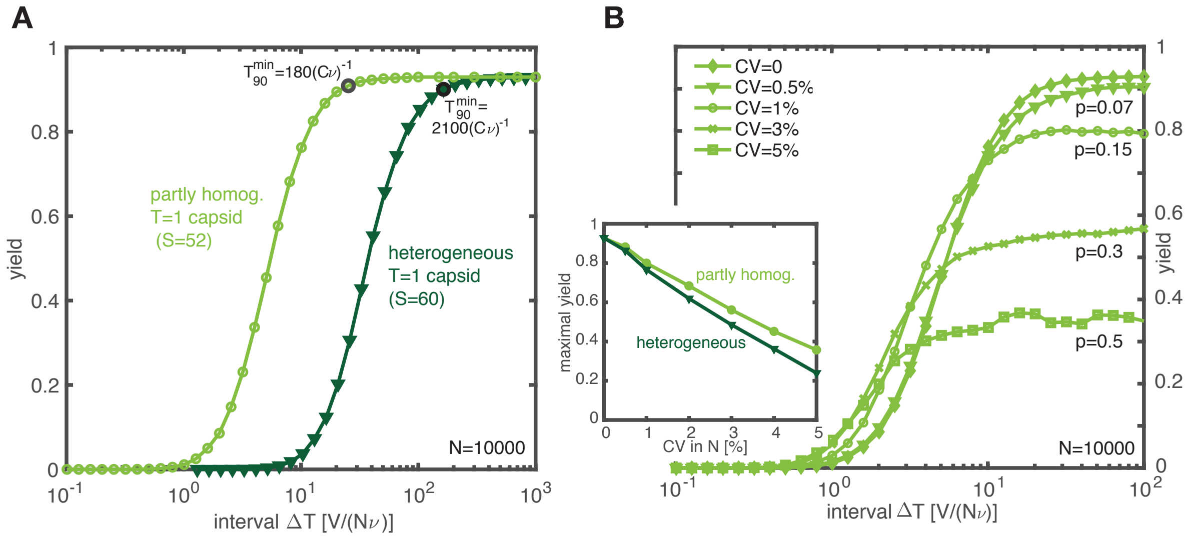


**Figure S7: Jis scenario for the T=1 capsid. A**, Final yield plotted against the interval $\Delta T$ between subsequent batches both for the partly homogeneous and the heterogeneous capsid (see Supplementary Fig. 6). Black circles indicate the position of the optimal interval $\Delta T_{\mathrm{opt}}$ and the corresponding minimal assembly time $T_{90}^{min}$. Simulations were performed for a maximal number of complete structures $N = 10^4$ and a fraction of resources that are distributed unevenly of $p = 0.07$ (cf. Methods and Eq. (33), which limits the yield to 93%. The partly homogeneous structure can be assembled faster than the heterogeneous structure, mainly because the binding speed is larger by a factor of 5 in the partly homogeneous capsid. **B**, Yield plotted against the interval $\Delta T$ for different levels of external noise in the particle numbers for the partly homogeneous capsid. For each species, the particle number from the protocol was perturbed independently with a specified coefficient of variation (CV := Gaussian standard deviation / mean). Inset shows the maximal yield for sufficiently large $\Delta T$ plotted against the coefficient of variation for the partly homogeneous and the heterogeneous structure. The fraction $p$ of resources that were distributed unevenly was chosen as follows: $p = 0.07$ for CV≤0.5%, $p = 0.15$ for CV=1%, $p = 0.2$ for CV=2%, $p = 0.3$ for CV=3%, $p = 0.36$ for CV=4% and $p = 0.5$ for CV=5%.
